# Cyclase associated actin cytoskeleton regulatory protein 2 (CAP2) organizes the actin cytoskeleton to influence the biomechanical properties of the ocular lens

**DOI:** 10.1101/2025.10.14.682085

**Authors:** Sepideh Cheheltani, Megan Coffin, Velia M. Fowler

**Author notes:** Corresponding Author: Velia M. Fowler, Ph.D., Department of Biological Sciences, University of Delaware, Newark, DE 19716.

## Abstract

Cyclase-associated actin cytoskeleton regulatory protein 2 (CAP2) is a conserved actin-binding protein that promotes actin filament (F-actin) turnover by disassembling ADF/cofilin-decorated filaments, supporting F-actin remodeling in differentiating cells and tissues. In the ocular lens, the actin cytoskeleton is critical for maintaining tissue biomechanical properties during fiber cell maturation. To assess CAP2’s role in the lens, we examined lens-specific CAP2 knockout (*CAP2^cKO^*) mice at cellular and tissue levels. *CAP2^cKO^* lenses were normal in size, shape, and transparency but exhibited increased stiffness under compression and enhanced recovery after load removal. While total actin levels and F-actin were unchanged, immunofluorescence revealed higher levels of Tropomodulin 1 (Tmod1, F-actin pointed-end capping), Tropomyosin3.5 (Tpm3.5, F-actin stabilizing), and T-plastin (F-actin bundling) in F-actin-rich membrane protrusions of mature fibers, while α-actinin-1 (F-actin cross-linking) was reduced. These findings suggest that CAP2 loss disrupts F-actin remodeling, promoting filament stabilization through Tpm3.5 binding, Tmod1 capping and T-plastin bundling. Consequently, F-actin networks become stiffer and more resilient, altering lens biomechanics. This study provides the first evidence that CAP2 regulates cell biomechanical properties in a non-muscle tissue through modulation of F-actin-associated proteins.

**Summary statement:** CAP2 loss alters the actin cytoskeleton through increased F-actin stabilization by Tmod1 and Tpm3.5, and filament bundling by T-plastin, leading to stiffer lenses.

## Introduction

Cyclase-associated Actin cytoskeleton regulatory Protein 2 (CAP2) is a conserved actin-regulatory protein (Peche et al., 2007; Rust et al., 2020) that modulates actin filament (F-actin) turnover by enhancing the depolymerization and severing of ADF/cofilin-bound F-actin from the pointed ends (Alimov et al., 2023; Kotila et al., 2019; Ono, 2013). CAP2 has been shown to play tissue-specific roles in organizing cytoskeletal architecture and supporting cellular functions (Field et al., 2015; Kosmas et al., 2015; Ono, 2013; Rust et al., 2020). In striated muscle, CAP2 localizes near the pointed ends of the thin filaments and is essential for proper myofibril assembly (Kepser et al., 2019); its deletion leads to disorganized actin networks, dilated cardiomyopathy, and impaired contractility (Colpan et al., 2021; Field et al., 2015; Peche et al., 2013). In non-muscle tissues such as the epidermis, CAP2 regulates cortical actin at cell junctions and promotes cell migration during wound healing (Kosmas et al., 2015; Peche et al., 2007). In neurons, CAP2 is a key regulator of actin dynamics and is essential for dendrite spine morphology and modulating synaptic plasticity (Kumar et al., 2016). However, whether CAP2 supports cytoskeletal and mechanical functions in highly specialized, non-contractile cells and tissues remains unclear. Here, we examine how CAP2 contributes to F-actin organization and biomechanics in the ocular lens, a transparent, deformable tissue that depends on a specialized F-actin network for cell differentiation and maturation.

The lens is a transparent spherical tissue in the eye’s anterior chamber, located behind the iris and in front of the vitreous humor (Lovicu and Robinson, 2004). It is composed of two types of cells, a monolayer of cuboidal epithelial cells on the anterior surface that overlays concentric layers of long, thin fiber cells extending from the anterior to the posterior poles, forming the vast majority of the lens mass (Bassnett and Mataic, 1997). The entire lens is encapsulated by a thin collagenous basement membrane, known as the capsule, which is produced by lens epithelial cells (Bassnett et al., 1999; Danysh and Duncan, 2009; Haddad and Bennett, 1988) (Fig. S1A). Lens function is dependent on two properties, transparency and biomechanics. The biomechanical properties of the lens, particularly its stiffness and elasticity, are essential for maintaining optical clarity and enabling accommodation (changes in lens shape to enable near and far focusing) (Cheng et al., 2019; Gupta et al., 2023). Age-related changes in lens stiffness contribute to the development of presbyopia (loss of lens ability to accommodate), underscoring the importance of understanding the molecular mechanisms that regulate lens mechanics. The eye lens presents a distinctive cellular context. Fiber cells are non-contractile and highly specialized, yet they rely on dramatic actin filament rearrangements during differentiation and maturation throughout life to maintain lens transparency. The lens is also unusual in that it continues to grow throughout life by the addition of new fiber cells at the equator (Fig. S1A) (Bassnett and Šikić, 2017). Disruptions in genes critical for lens development or fiber cell organization often lead to smaller lenses in adult mice, demonstrating how early defects can have lasting effects on lens size (Graw, 2019; Li et al., 2024). For instance, mutations in Connexin 50, Aquaporin 0, or lens intrinsic membrane protein-2 (Lim2) result in a smaller lens phenotype (Gu et al., 2019; Shiels et al., 2007), emphasizing the importance of precise cellular organization during development for normal lens growth and biomechanics.

During maturation, lens fiber cells develop specialized paddles and interlocking membrane interdigitations enriched in F-actin networks with diverse associated actin binding proteins (ABPs) (Bassnett and Šikić, 2017; Cheng et al., 2016c; Šikić et al., 2015) (Fig. S1B). This complex architecture is arranged by a host of ABPs that regulate filament assembly, turnover, and organization (Cheng et al., 2017). Previous research investigating the roles of ABPs in lens actin networks and biomechanical properties revealed that the reduction or deletion of F-actin-stabilizing proteins, tropomyosin isoform 3.5 (Tpm3.5) and tropomodulin 1 (Tmod1), leads to F-actin network rearrangements and altered lens biomechanical properties with reduced lens stiffness (Cheng et al., 2018; Cheng et al., 2016c; Fischer et al., 2000; Gokhin et al., 2012). Tpms and Tmods are actin-stabilizing proteins that protect F-actin from ADF/Cofilin-mediated disassembly and depolymerization in a variety of cells (Gunning et al., 2015; Jansen and Goode, 2019; Ono and Ono, 2002). However, the role of actin disassembly proteins, such as ADF/Cofilin and CAP, in the turnover and rearrangement of the lens actin cytoskeleton network is unknown (Bamburg and Bernstein, 2010; Cheng et al., 2017). A key question and the central focus of this study is what role CAP2 may play in regulating actin networks in the lens and how it interfaces with other ABPs to maintain the unique architecture of fiber cells.

Notably, CAP2-knockout mice develop microphthalmia (Field et al., 2015), indicating that CAP2 may be important for normal lens growth or architecture. Here, we have investigated how CAP2 fits into the network of ABPs in the lens. We ask whether CAP2 regulates actin filament remodeling in fiber cells and cooperates or antagonizes with known lens ABPs to assemble and maintain the cytoskeletal F-actin network. By investigating the role of CAP2 within the broader ecosystem of filament regulators, we aim to understand how the actin cytoskeleton of the lens is affected in the absence of a key actin regulatory protein and provide insight into the general principles by which tissue-specific actin networks are regulated to support specialized cellular functions. In the following, we present a focused analysis of CAP2’s role in the lens, detailing its expression and functional impact on fiber cell organization and lens biomechanics, thereby shedding light on how this actin regulator contributes to the complex requirements of lens flexibility.

## Results

### CAP2 protein is absent in CAP2^cKO^ mice lens

The survival rate for CAP2 global KO mice is very low, with the few that survive after birth exhibiting multiple defects, including severe cardiomyopathy, and typically dying before ten weeks of age (Field et al., 2015). Therefore, to investigate CAP2’s function in the adult mouse lens and to circumvent potential systemic effects of global CAP2 deletion, we utilized a conditional knockout strategy to delete CAP2 in lens epithelium and fiber cells using the Cre-Lox system (Fig. S2). Specifically, we used the *MLR10-Cre* transgenic mouse, which induces CRE expression in lens epithelial and fiber cells at embryonic day 10.5 (E10.5) (Zhao et al., 2004), to generate a lens-specific CAP2 knockout mouse (Lam et al., 2019). RT-qPCR and western blot analysis confirmed the absence of *Cap2* mRNA and protein in *CAP2^fl/fl^ ^Tg^ ^(Cryaa-Cre)10Mlr^* (Hereafter referred to as *CAP2^cKO^*) lens samples (Fig. 1A, C). To confirm that the deletion of CAP2 is lens-specific, we immunostained 8-week-old lens frozen sections in the equatorial and sagittal orientations with anti-CAP2 rabbit antibody. Immunostaining of mouse eye sections revealed that CAP2 is expressed in the lens, cornea, and retina (Fig. S3A). Low-magnification confocal images revealed that the CAP2 signal is absent in *CAP2^cKO^* lens, in contrast to the robust CAP2 staining observed in non-lens tissues such as the ciliary body (Fig. S3B). Additionally, although we focused on CAP2, vertebrates express both *Cap1* and *Cap2* genes (Rust et al., 2020). To assess potential compensatory mechanisms, we examined whether *Cap1* expression is altered in *CAP2^cKO^* lenses. However, we detected no change in *Cap1* mRNA levels in the absence of *Cap2* (Fig. 1B). According to the iSyTE database of lens gene expression (Lachke et al., 2012), *Cap2* is more abundantly expressed and highly enriched in the lens compared to *Cap1*. Thus, it is unlikely that *Cap1* expression compensates for the loss of CAP2 in knockout lenses.

**Figure 1.**
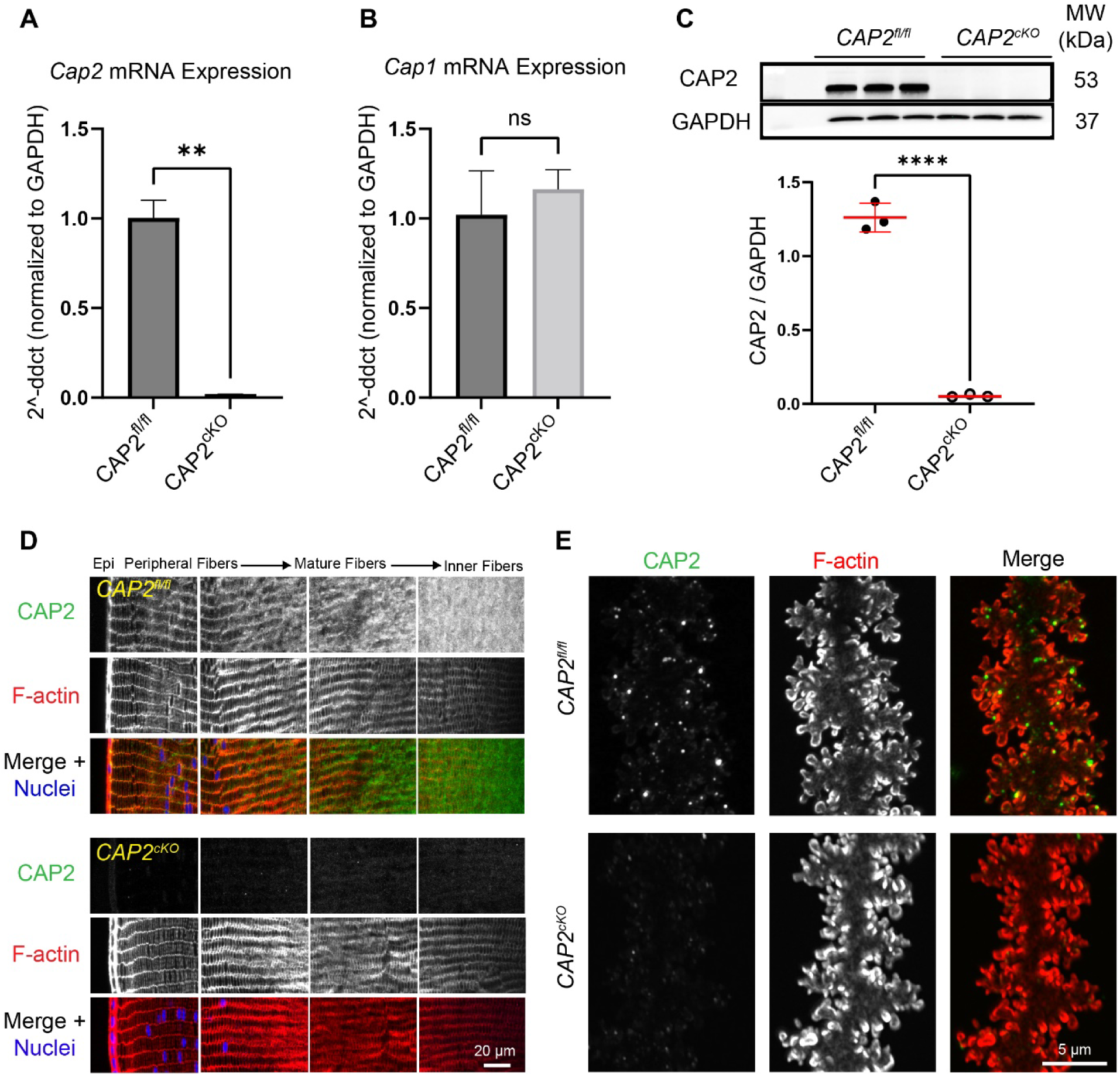
CAP2 is not present in *CAP2^cKO^* mouse lenses. (A) RT-qPCR from three *CAP2^cKO^* and *CAP2^fl/fl^*mouse lenses reveals a significant decrease in transcript levels in the KO samples. (B) RT-qPCR from three *CAP2^cKO^* and *CAP2^fl/fl^* mouse lenses reveals no significant changes in mRNA levels of Cap1 in lenses. GADPH was used as a housekeeping gene. (C) Western blots of *CAP2^cKO^* and *CAP2^fl/fl^* lenses reveal the absence of CAP2 protein in KO samples. GAPDH is used to normalize protein levels. Plots reflect mean ± s.d. of n=3 independent samples per genotype. **P <0.01; ****P <0.0001. (D) Immunostaining of the equatorial cryosections of 8-week-old *CAP2^fl/fl^* and *CAP2^cKO^*lenses stained with anti-CAP2 rabbit polyclonal antibody (green), rhodamine phalloidin for F-actin (red), and Hoechst for nuclei (blue). In control lens fiber cells, CAP2 staining appears to be enriched with F-actin on the broad sides, at the vertices, and diffuse in the cytoplasm. (E) Immunostaining of single fiber cells from 8-week-old *CAP2^cKO^* and *CAP2^fl/fl^*lenses for CAP2 (green) and F-actin (red). Images are maximum intensity projections. Scale bars, 20 µm (D), 5 µm (E).

### CAP2 is present in puncta near F-actin-rich protrusions of lens fiber cells

To study the role of CAP2 in the lens actin cytoskeleton, we first examined the expression and localization of CAP2 protein in the mouse lens (Fig. 1D). In equatorial cryosections, CAP2 labeling is prominent in lens fiber cells with minimal signal in lens epithelial cells. In fiber cells, CAP2 staining appears enriched with F-actin on the broad sides and at the vertices, as well as diffuse in the cytoplasm. Additionally, CAP2 is localized with F-actin near the membrane in both differentiating and maturing fiber cells, but is more abundant in the cytoplasm of inner cortical fiber cells (Fig. 1D). Taking a closer look at the subcellular localization of CAP2 in the mature region of fiber cells revealed that CAP2 appears as puncta near the F-actin-rich protrusions (Fig. 1E). Together, these data show that CAP2 is predominantly cytoplasmic, with puncta near F-actin-rich membrane protrusions, suggesting that a portion of CAP2 associates with the actin cytoskeleton in lens fiber cells.

### CAP2^cKO^ lenses are normal in size and shape but exhibit increased stiffness

To evaluate lens morphology in *CAP2^cKO^* mice, we conducted a comprehensive examination of whole lens morphology by analyzing freshly dissected lenses from 8-week-old *CAP2^cKO^* and control mice (*CAP2^fl/fl^*). Our initial image analysis revealed that the lenses from *CAP2^cKO^* mice were transparent, with no apparent opacities (Fig. 2A). Morphometric analyses further revealed no significant alterations in either lens total volume or shape (aspect ratio) in *CAP2^cKO^*lenses compared to control littermates for either male or female mice (Fig. 2B). Additionally, the nuclear volume and nuclear fraction remained unchanged in the mutant lenses (Fig. 2C). These findings suggest that CAP2 does not directly regulate lens development and growth.

**Figure 2.**
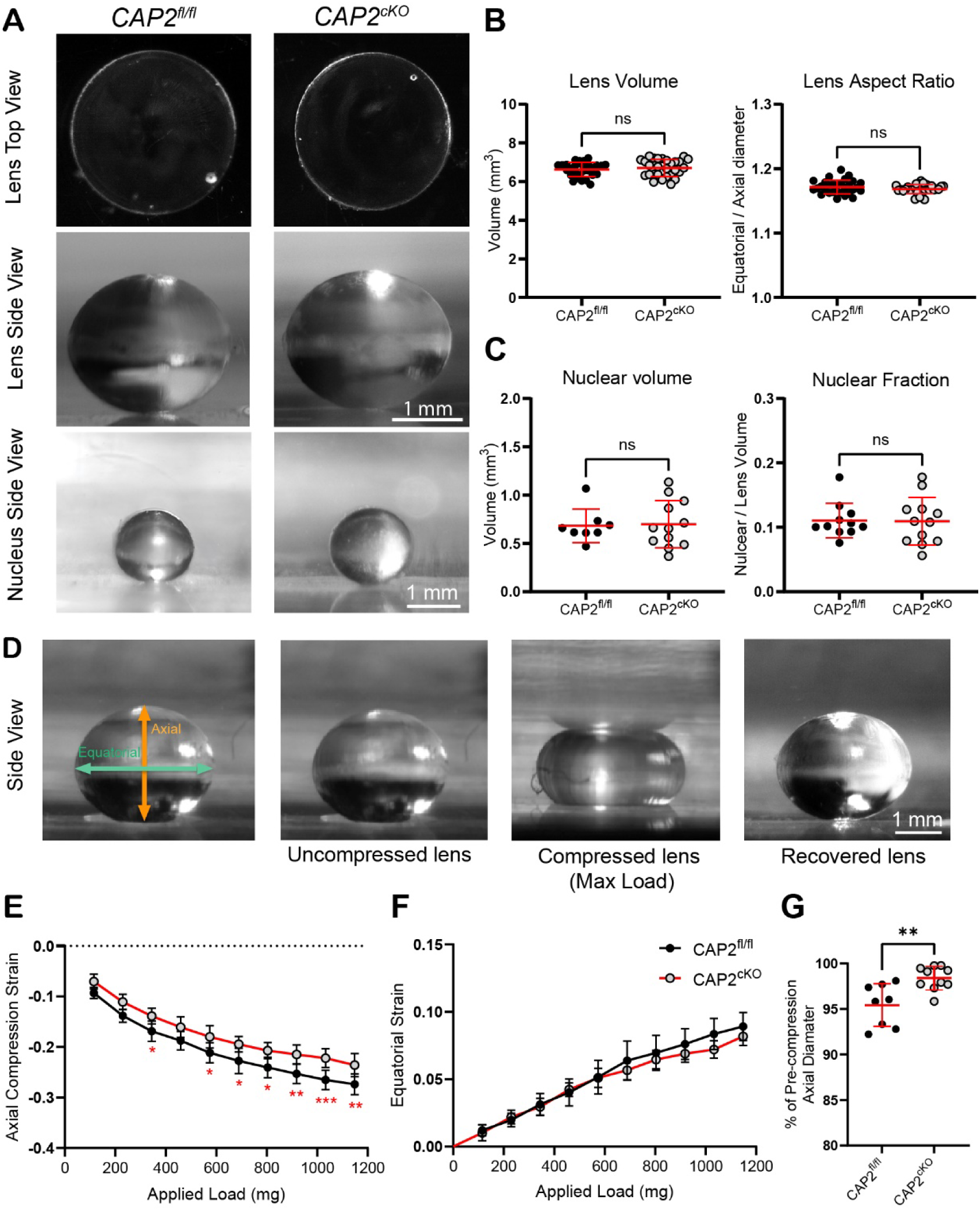
*CAP2^cKO^* lenses have normal morphology but are stiffer and more resilient. (A) images of freshly dissected 8-week-old *CAP2^cKO^*and *CAP2^fl/fl^* lenses. All lenses were transparent without obvious opacities. (B) Whole lens volume and aspect ratio (equatorial to axial diameter ratio) from 8-week-old control and mutant mice show no significant differences in whole lens size and shape. (C) Nuclear fraction and volume show no significant difference in nuclear volume in mutant lenses. Plots show mean ± s.d. of 12-15 lenses from at least seven biological replicates per genotype. (D) Side view images of uncompressed, compressed, and recovered lenses. (E, F) Plots of axial and equatorial strain (d−d0)/d0 show increased axial strain in *CAP2^cKO^* lenses, especially at high loads. (G) *CAP2^cKO^*lenses had increased resilience, calculated as the ratio of the pre-compression to post-compression axial diameter. Plots reflect the mean ± s.d. of 8 lenses from at least five mice per genotype. \**P* <0.05; \*\**P* < 0.01; \*\*\**P* < 0.001. Scale bars, 1 mm.

To evaluate whether the absence of CAP2 alters the biomechanical properties of the lens, we conducted a coverslip compression assay to measure lens stiffness and the ability to recover (Cheng et al., 2016a) (Fig. 2D). Our analysis revealed a significant decrease in axial compressive strain in *CAP2^cKO^*lenses, particularly at higher loads, indicating that *CAP2^cKO^*lenses are stiffer than control lenses under compression (Fig. 2E, F). Additionally, we evaluated lens shape recovery after load removal by calculating the ratio of the pre- to post-loading axial diameter, referred to as resilience. Our findings demonstrated a significant improvement in recovery in *CAP2^cKO^* lenses, with a mean recovery to 98.39 ± 1.28% (mean ± SD) of the pre-loading axial diameter, as compared to 95.42 ± 2.337% for the control lenses (Fig. 2G). These results are unexpected, as previous work on mice with reduced levels of ABPs, such as Tpm3.5, Tmod1, and intermediate filament protein, CP49, led to softer lenses with decreased resilience, specifically in Tpm3.5-deficient lenses (Cheng et al., 2018; Gokhin et al., 2012).

Studies of decapsulated lenses have demonstrated that the capsule contributes to lens shape retention and resilience (Mekonnen et al., 2023; Wilde et al., 2012). To assess whether changes in capsule thickness might contribute to the increased stiffness and enhanced recovery of *CAP2^cKO^* lenses, we measured anterior capsule thickness in both *CAP2^cKO^* and *CAP2^fl/fl^* lenses (Fig. S4A-C). Image analysis revealed no significant differences between genotypes (Fig. S4C), indicating that capsule thickness is unaffected by CAP2 deletion, which is consistent with the minimal expression of CAP2 in the lens epithelium.

Since changes in lens biomechanics can sometimes arise from alterations in fiber cell organization (Cheng et al., 2016c; Gu et al., 2019; Maddala et al., 2011), we next examined lens fiber cell organization in CAP2^cKO^ lenses to determine if the altered mechanics was accompanied by structural defects. Confocal imaging of lens cryosections showed that fiber cells in CAP2^cKO^ lenses maintained their elongated shape and typical hexagonal packing in cross-section (Fig. S4D). The characteristic alignment of fiber cells into radial cell columns and the formation of anterior and posterior sutures also appeared normal in cKO lenses (data not shown). Minor deviations in fiber cell packing were observed in cKO lenses near the equator (Fig. S4D, E), but these changes were subtle and unlikely to account for the observed biomechanical phenotype, since other genetically-modified lenses with extensive fiber cell hexagonal packing defects have no changes in biomechanical stiffness (Islam et al., 2023).

### Loss of CAP2 does not affect the ratio of F-actin: G-actin in lens fiber cells

Based on the ability of CAP2 to promote actin filament disassembly (Colpan et al., 2021; Kosmas et al., 2015; Peche et al., 2007), we hypothesized that deleting CAP2 in the lens might similarly alter actin polymerization and affect the proportion of Filamentous to Globular actin (F- to G-actin). To test this, we performed complementary biochemical (Fig. 3A-E) and immunostaining approaches (Fig. 3F-I). The total actin level was unchanged in *CAP2^cKO^* lenses, based on western blotting (Fig. 3B). Subcellular fractionation to isolate lens cytosol (supernatant) and membrane and cytoskeleton fractions (pellets), followed by western blotting, demonstrated that ∼ 69% of actin is present in the lens cytosol in the *CAP2^fl/fl^* lenses, as shown previously (Woo and Fowler, 1994), with no increase in cytosolic levels of actin in the *CAP2^c^*^KO^ lenses (Fig. 3B, D). We also observed that ∼93% of CAP2 protein is cytosolic in control lenses (Fig. 3B, E), consistent with previous reports showing CAP2 distributed diffusely in the cytoplasm of mouse embryonic fibroblasts (Kepser et al., 2021) and cardiomyocytes (Colpan et al., 2021), in addition to its enrichment at the pointed ends of actin filaments in cardiomyocytes (Colpan et al., 2021). Therefore, we conclude that the absence of CAP2 does not significantly influence the overall F to G-actin ratio within lens fiber cells.

**Figure 3.**
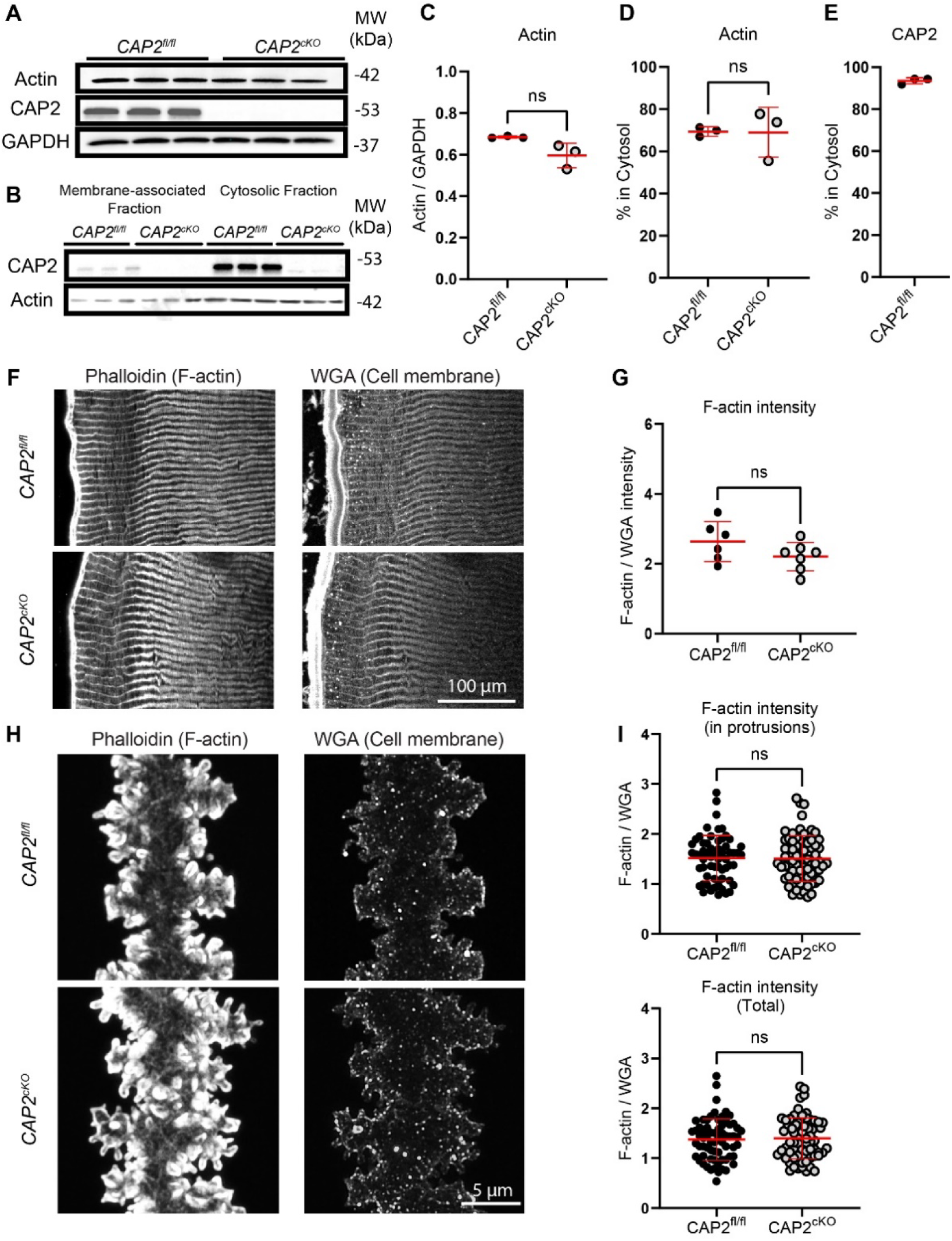
The ratio between cytosolic and membrane-associated fractions of actin is similar in *CAP2^cKO^* and control lenses. Western blot of lens total proteins (A) and cytoplasmic and membrane fractions (B). The percentage of each protein in the cytosol was calculated by dividing the cytosol band intensity by the intensity of the cytosol band plus the membrane band. There is no change in total actin levels or (C) the percentage of actin and CAP2 in the cytosol in mutant lenses (D, E). Plots reflect mean ± s.d. of 3 independent protein samples per genotype. (F) Immunostaining of lens cryosections (F) and single fiber cells (H) with rhodamine phalloidin for F-actin and WGA for the cell membrane. The signal intensity of F-actin, normalized to WGA, is not different between mutant and control lenses (G, I, J). Plots reflect mean ± s.d. of 6 sections from 6 mice per genotype (G) and a total of 70 individual fibers from 9 mice per genotype (I, J). Scale bars, 100 µm (F), 5 µm (H).

We also investigated F-actin levels in control (*CAP2^fl/fl^*) and *CAP2^c^*^KO^ fiber cells by staining F-actin in lens cryosections with rhodamine-phalloidin, using Wheat Germ Agglutinin (WGA) as a marker for fiber cell membranes (Kistler et al., 1986). This showed that loss of CAP2 had no effect on the fluorescent intensity of F-actin with respect to WGA (Fig. 3F, G). Next, we considered whether the loss of CAP2 might lead to altered abundance of F-actin in different lens regions containing young, maturing, and older lens fiber cells. However, the ratio of F-actin to WGA revealed no significant differences in F-actin fluorescent intensity between the *CAP2^cKO^* lens and the control in fiber cells at different depths (data not shown).

We next considered the possibility that changes in actin polymerization in specific F-actin-rich structures, such as membrane protrusions, could underlie the increased stiffness. The F-actin-rich lens fiber cell membrane protrusions have been demonstrated to play a crucial role in determining the lens’s biomechanical properties (Cheng et al., 2018; Gokhin et al., 2012). Recently, we showed that compressive loading of lenses induces significant curvature in cortical fiber bundles, accompanied by reversible distortion of membrane paddles and effacement of the small membrane protrusions (Cheheltani et al., 2025b). To investigate whether loss of CAP2 affects F-actin levels within these protrusions, we quantified F-actin signal intensity specifically within the membrane protrusion domains of single mature fiber cells (Fig. 3H). F-actin abundance within these protrusions was not significantly different from that of controls (Fig. 3I), suggesting that altered actin network organization, rather than filament polymerization and accumulation, could be the underlying mechanism causing the stiffer lens phenotype.

### Loss of CAP2 alters the levels of some actin-associated proteins in the lens

Prior studies revealed that ABPs modulate lens biomechanics by altering cytoskeletal networks and F-actin-stabilizing proteins in fiber cells. For example, the absence of Tmod1 destabilizes the spectrin-F-actin network, leading to softer lenses (Cheng et al., 2016c). Likewise, Tpm3.5 deficiency results in dissociation of Tmod1 from F-actin at the membrane, as well as altered distribution of spectrin and reduced levels of T-plastin, leading to decreased lens stiffness under high loads (Cheng et al., 2018). Therefore, we hypothesized that the increased lens stiffness observed in the absence of CAP2 might stem from the opposite effect. In other words, loss of CAP2 could trigger the upregulation or redistribution of F-actin stabilizers and membrane linkage proteins that reinforce the cortical cytoskeleton, thereby increasing tissue stiffness. To test this idea, we examined whether the absence of CAP2 alters the levels or localization of key F-actin stabilizing and crosslinking proteins in the lens.

First, to determine whether the absence of CAP2 affects the abundance of other cytoskeletal proteins, we assessed the total protein levels of key ABPs in whole-lens extracts from *CAP2^cKO^* and control mice. Western blot analysis revealed several notable changes (Fig. 4). Tmod1, a pointed-end capping protein that stabilizes actin filaments, was upregulated in CAP2-deficient lenses. In contrast, α-actinin-1, an anti-parallel F-actin cross-linker, was significantly reduced. T-plastin, which bundles parallel F-actins, was unchanged between genotypes, while ezrin, a membrane-cytoskeleton linker associated with F-actin-rich protrusions, was elevated in the *CAP2^cKO^* lenses. Notably, the levels of Tpm3.5 and the core membrane scaffold protein β2-spectrin remained unchanged, suggesting that loss of CAP2 deletion does not broadly perturb all actin cytoskeletal components.

**Figure 4.**
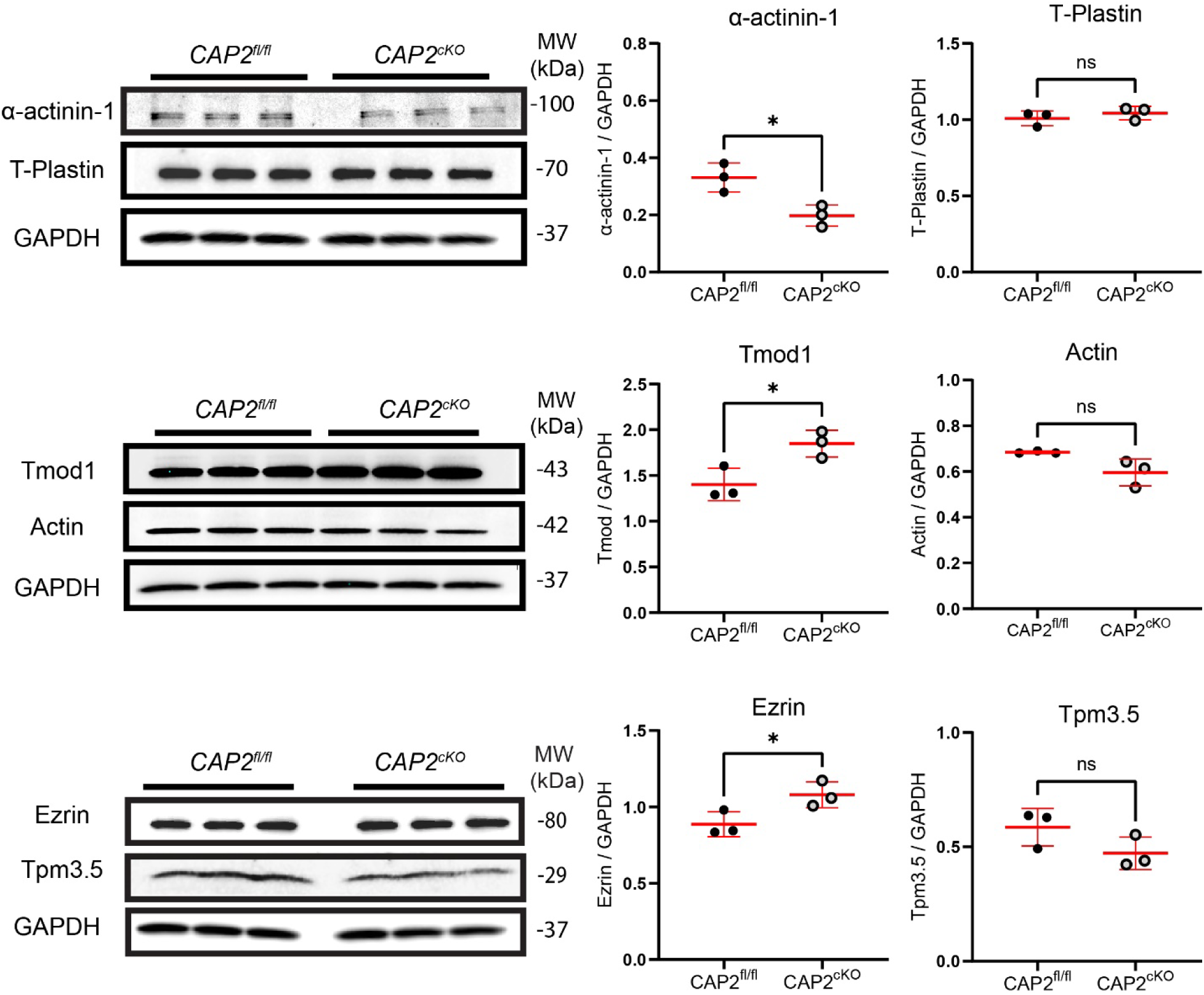
Deletion of CAP2 alters levels of some actin-associated proteins: α-actinin-1, Tmod1, and ezrin. Western blots of actin-associated proteins from 8-week-old whole lenses. All protein levels were normalized to GAPDH. Total levels of α-actinin-1 protein are decreased in *CAP2^cKO^*lenses, while T-plastin levels are unchanged. Tmod1, an actin-stabilizing protein, is increased in the mutant lenses, while the level of actin and Tpm3.5 is unaffected. The level of ezrin, an actin cytoskeleton to membrane linking protein, is increased in mutant lenses. Plots show the mean ± s.d. of 3 independent protein samples per genotype. \**P* <0.05.

### F-actin-stabilizing proteins are increased in CAP2^cKO^ lenses

CAP2 regulates filament turnover at pointed ends (Colpan et al., 2021; Iwanski et al., 2021; Kotila et al., 2019); therefore, we investigated whether its deletion affects the distribution of Tmod1, the pointed-end capping protein in fiber cells. We first examined the fluorescent intensity of Tmod1 in lens equatorial cryosections at fiber cell regions extending from the epithelium to the inner fiber cells (Fig. S5A). Quantification of Tmod1/F-actin intensity revealed that Tmod1 fluorescent intensity is significantly higher in *CAP2^cKO^* lenses compared to control lenses, especially in the region spanning outer cortical and mature fiber cells (0-160 μm inwards from the epithelial cells) (Fig. S4B). Further, closer examination of Tmod1 localization in individual mature fiber cells revealed that Tmod1 is associated with puncta in the F-actin membrane protrusions of mature fiber cells in control and *CAP2^cKO^* lenses (Fig. 5A). We quantified the fluorescent intensity of the Tmod1 puncta in the protrusions and normalized it to F-actin intensity for individual mature fiber cells from control and *CAP2^cKO^* lenses. This revealed that in *CAP2^cKO^* lenses, the total intensity of Tmod1/F-actin, and the intensity of Tmod1/F-actin in the membrane protrusions of mature fibers is higher (Fig. 5A). The increased staining intensity of Tmod1 with respect to F-actin in *CAP2^cKO^* lenses suggests that loss of CAP2 leads to increases in Tmod1-capped F-actin in lens fiber cells.

**Figure 5.**
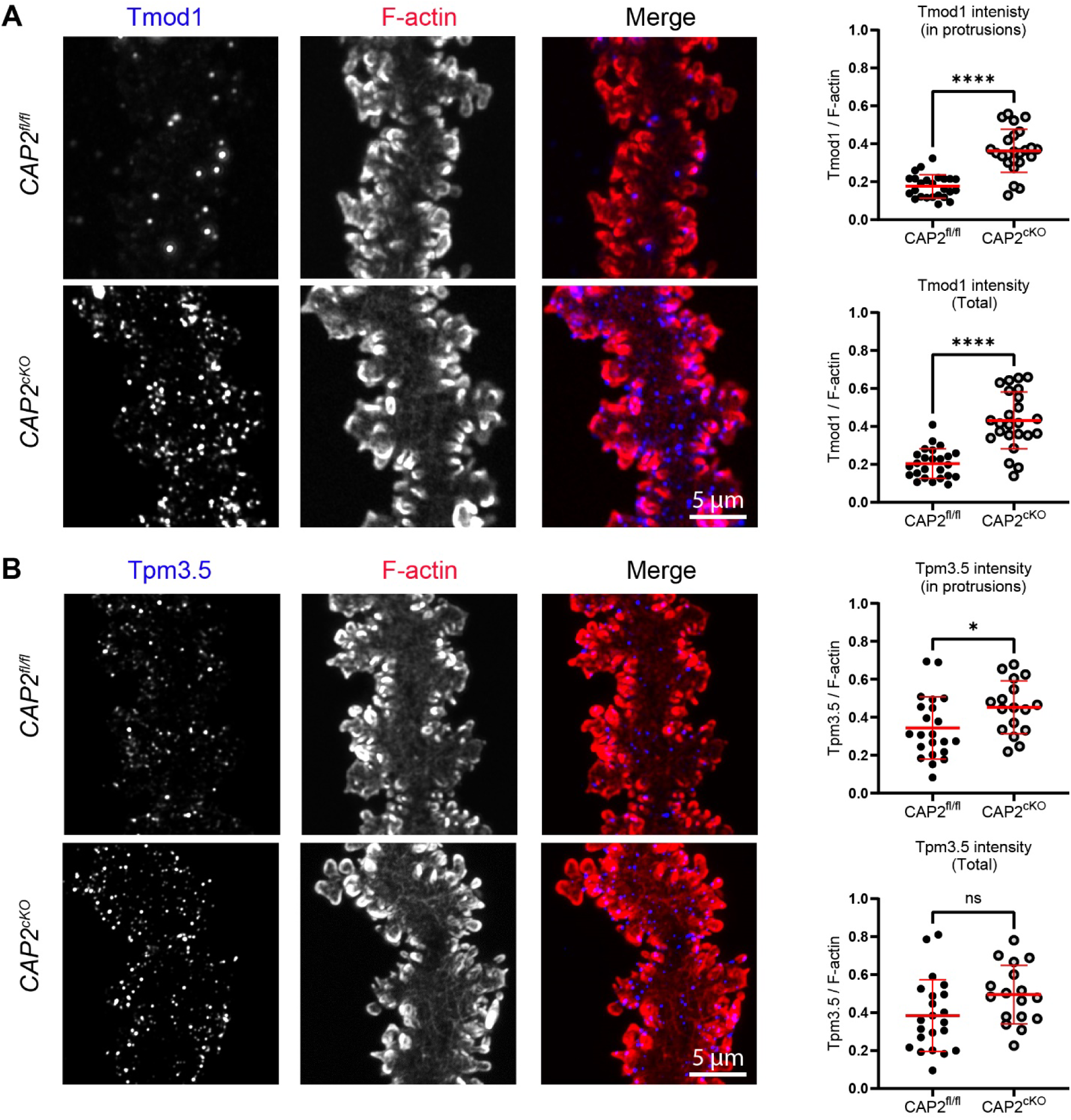
Tmod1 and Tpm3.5 are increased in *CAP2^cKO^* lenses. (A) Immunostaining of single mature fiber cells from 6-week-old *CAP2^cKO^* and *CAP2^fl/fl^* lenses for Tmod1 (blue) and F-actin (red). Images are maximum intensity projections. Tmod1 is localized in puncta near F-actin-rich membrane protrusions in control lenses and the signal intensity is markedly increased in both protrusions and the whole fiber cell in CAP2^cKO^ fibers. (B) Immunostaining of a single mature fiber cell for Tpm3.5 (blue) and F-actin (red). (B) Tpm3.5 is enriched in puncta near the F-actin-rich membrane protrusions. While its total intensity is unchanged, its intensity at protrusions is significantly increased in *CAP2^cKO^* lens fibers. Scale bars, 5 µm. Plots represent the mean ± s.d. of 20-30 individual fibers from 6 lenses of 3 different mice per genotype. \**P* < 0.05; \*\*\*\**P* <0.0001.

Since Tmod1 and Tpms are essential F-actin-stabilizing proteins that interact with one another, and previous studies showed their importance in lens biomechanical properties (Cheng et al., 2018; Gokhin et al., 2012), we also evaluated the localization and abundance of the major Tpm isoform in lens fiber cells, Tpm3.5 (Cheng et al., 2018; Cheng et al., 2016c). Consistent with enhanced pointed-end stability, we detected an increase in Tpm3.5 localization associated with F-actin in or near the membrane protrusions of *CAP2^cKO^* mature fiber cells (Fig. 5B). This enrichment of Tpm3.5 with respect to F-actin in the absence of CAP2 implies that more actin filaments are decorated with Tpm3.5, and available for high affinity capping by Tmod1 near the membrane of cKO fibers.

### CAP2 absence disrupts α-actinin-1-linked actin networks and increases T-plastin-linked F-actin bundles at fiber cell membranes

In mouse lenses lacking F-actin-stabilizing proteins such as Tmod1 or Tpm3.5, the F-actin organization in membrane protrusions is disrupted, with aberrant distributions of actin cross-linking proteins such as α-actinin-1 and T-plastin (Cheng et al., 2018; Cheng et al., 2016c). For example, the softer Tpm3.5-deficient lenses exhibit increased prevalence of α-actinin-1 networks (Cheng et al., 2018), while softer Tmod1-knockout lenses display expanded, nearly continuous α-actinin-1 localization along the fiber cell membrane, distinct from the punctate pattern observed in controls (Cheng et al., 2016c). Given these results, and the observed increase in lens stiffness in *CAP2^cKO^* lenses, we investigated whether decreases in α-actinin-1 might be observed in the mature single fiber cells without CAP2 (Fig. 6). In control lens fiber cells, prominent α-actinin-1 puncta are found on the side, tips, and at the base of the small F-actin-rich membrane protrusions (Fig. 6A), while in the fiber cells from *CAP2^cKO^* lenses, the α-actinin-1 puncta relative to F-actin are significantly reduced (Fig. 6A). A decrease in α-actinin-1 intensity relative to F-actin is also observed in younger, peripheral fiber cells, based on immunostaining of equatorial cryosections (Fig. S6). These results suggest that loss of CAP2 leads to reductions in α-actinin-1-linked F-actin networks in peripheral and mature fiber cells.

**Figure 6.**
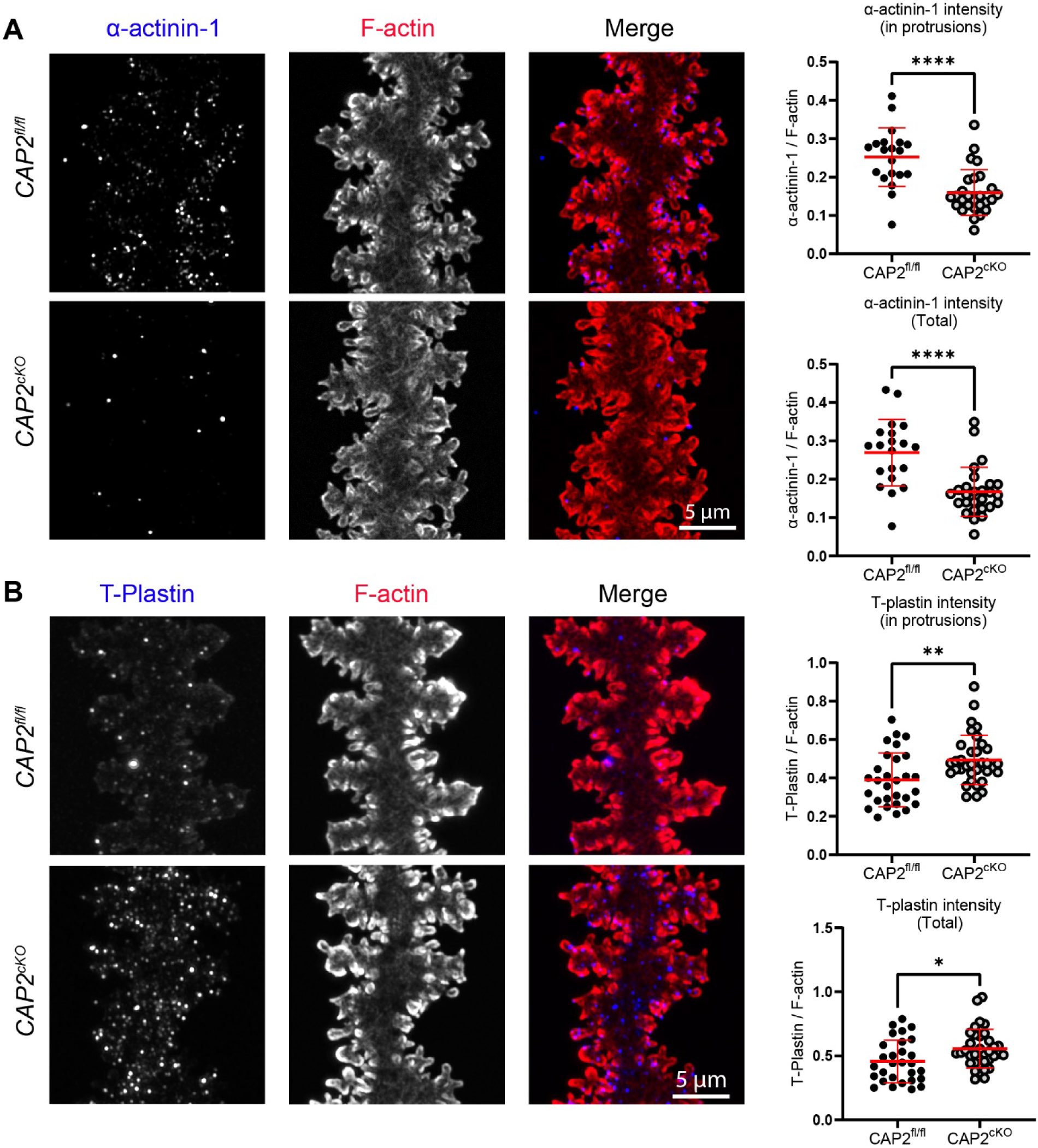
The α-actinin-1-F-actin network is decreased while T-plastin-F-actin network is increased in *CAP2^cKO^* lens fibers. (A) Immunostaining of single mature fiber cells from 6-week-old *CAP2^fl/fl^* and *CAP2^cKO^* lenses for α-actinin-1 (blue) and F-actin (red). Images are maximum intensity projections. In the control lens fiber, α-actinin-1 is localized to the base and tips of protrusions. In the mutant fibers, α-actinin-1 levels are significantly reduced in both protrusions and overall. (B) Immunostaining of single mature fiber cells from 6-week-old lenses for T-plastin (blue) and F-actin (red). T-plastin is mostly localized in small puncta at the base of the paddles of the control lens fiber. While the localization of T-plastin does not appear affected in mutant lenses, the intensity of T-plastin in the protrusion region and overall is significantly greater in *CAP2^cKO^* lenses. Scale bars, 5 µm. Plots reflect the mean ± s.d. of 20-30 individual fibers from 6 lenses of 3 different mice per genotype. \**P* < 0.05; \*\**P* <0.0; \*\*\*\**P* <0.0001

In contrast to α-actinin-1 networks, T-plastin networks were reduced in Tpm3.5-deficient lenses (Cheng et al., 2018). Therefore, we next examined T-plastin to determine whether its localization or expression was affected in CAP2-deficient lenses. In control lens fiber cells, sparse T-plastin puncta are associated with F-actin in membrane paddles and protrusions (Fig. 6B). In contrast, in lens fiber cells from *CAP2^cKO^* lenses, the T-plastin puncta appear more abundant and their intensity relative to F-actin is significantly increased (Fig. 6B). This suggests that loss of CAP2 leads to expansion of F-actin bundles associated with T-plastin at the membrane protrusions of mature lens fibers.

### Loss of CAP2 leads to increased Ezrin, while β2-Spectrin localization remains unchanged

Since loss of Tmod1 leads to softer lenses and is associated with disrupted spectrin-actin networks (Cheng et al., 2016c), we hypothesized that the increased stiffness in CAP2-deficient lenses may involve the opposite effect, reinforcement of actin-membrane linkages. We therefore tested whether the levels or localization of key membrane-cytoskeleton linkers in the lens, such as ezrin and β2-spectrin, are altered in the absence of CAP2. Consistent with the increase in total ezrin levels in western blot of *CAP2^cKO^* lenses shown above (Fig. 4), immunostaining showed an increase in ezrin puncta intensity in mature single lens fiber cells (Fig. 7A). In contrast, the localization and abundance of β2-spectrin, an essential scaffolding protein of the membrane-associated spectrin-actin cytoskeleton, remains unchanged in CAP2-deficient lenses (Fig. 7B). This suggests that the core spectrin-actin membrane skeleton is preserved, but specific membrane actin linkers, such as ezrin, may be selectively upregulated to compensate for the other alterations in F-actin network organization. Together, these protein-level changes reflect targeted reorganization of the actin cytoskeletal network in CAP2-deficient lenses, particularly in the F-actin-enriched membrane protrusion regions of mature fiber cells.

**Figure 7.**
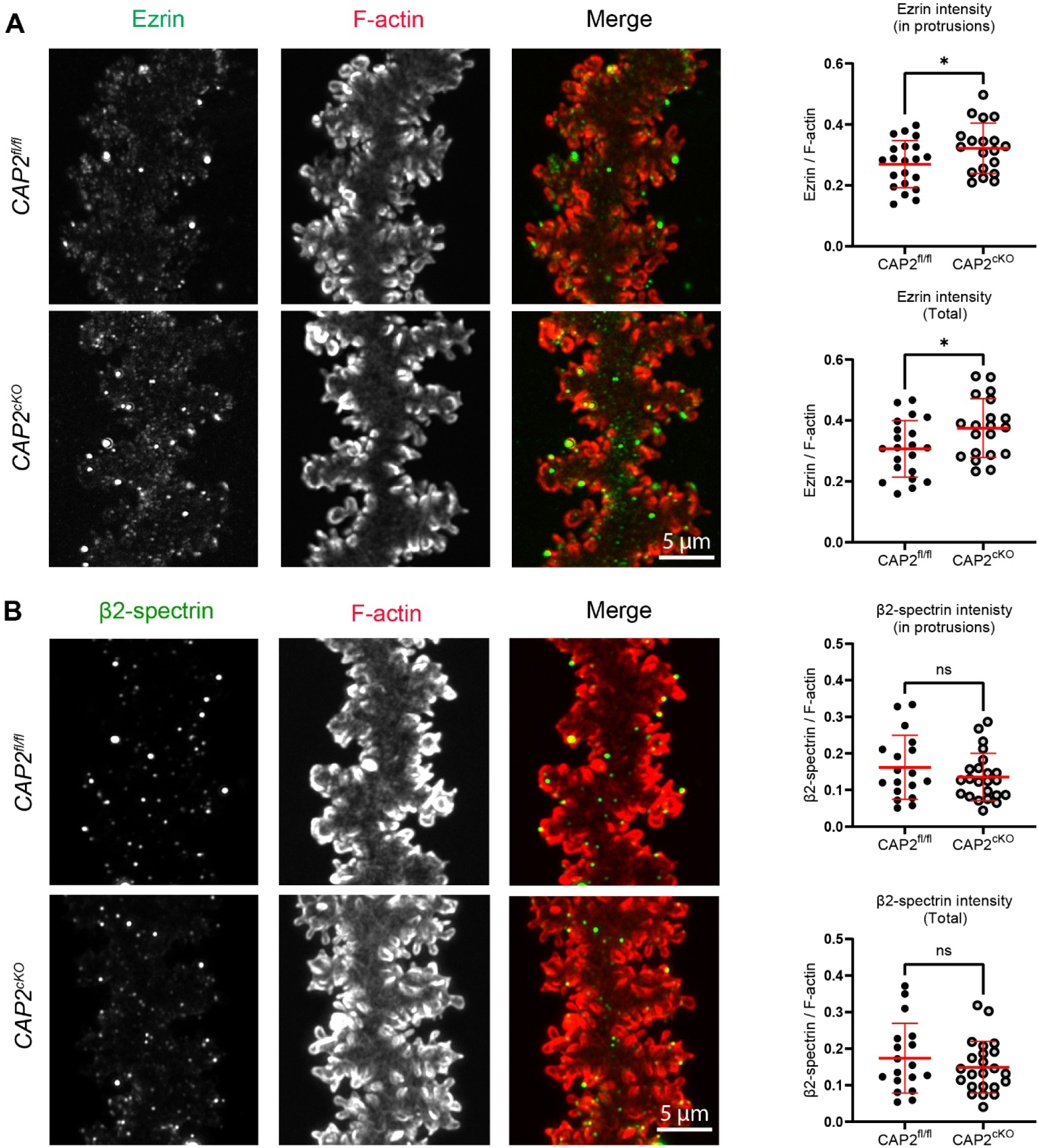
Ezrin is increased in *CAP2^cKO^* lenses. (A) Immunostaining of single mature fiber cells from 6-week-old *CAP2^fl/fl^* and *CAP2^cKO^* lenses for ezrin (green) and F-actin (red). Images are maximum intensity projections. In the control lens fiber, ezrin is enriched near the membrane protrusions. In the KO lens fiber, the staining is significantly increased. (B) Immunostaining of single mature fiber cells from 6-week-old lenses for β2-spectrin (green) and F-actin (red). β2-spectrin is mostly localized at the base of the paddles and the tip of some small protrusions in the control lens fiber. The localization and abundance of β2-spectrin in *CAP2^cKO^* lenses are unchanged. Scale bars, 5 µm. Plots represent the mean ± s.d. of 20-30 individual fibers from 6 lenses of 3 different mice per genotype. \**P* < 0.05.

## Discussion

The biomechanical properties of the ocular lens are dependent on precise regulation of its cytoskeletal architecture, particularly within the F-actin-rich cortical regions of fiber cells (Cheng et al., 2018; Cheng et al., 2017; Gokhin et al., 2012). F-actin stability, crosslinking, and membrane tethering all contribute to the mouse lens’s ability to deform and recover in response to mechanical compression and load release (Cheng et al., 2018; Gokhin et al., 2012). Our current study identifies CAP2 as one of the key regulators of these processes, acting locally near the membrane of mature fiber cells to influence the structural organization of the F-actin cytoskeleton and mechanical properties of the lens (Fig 8).

**Figure 8.**
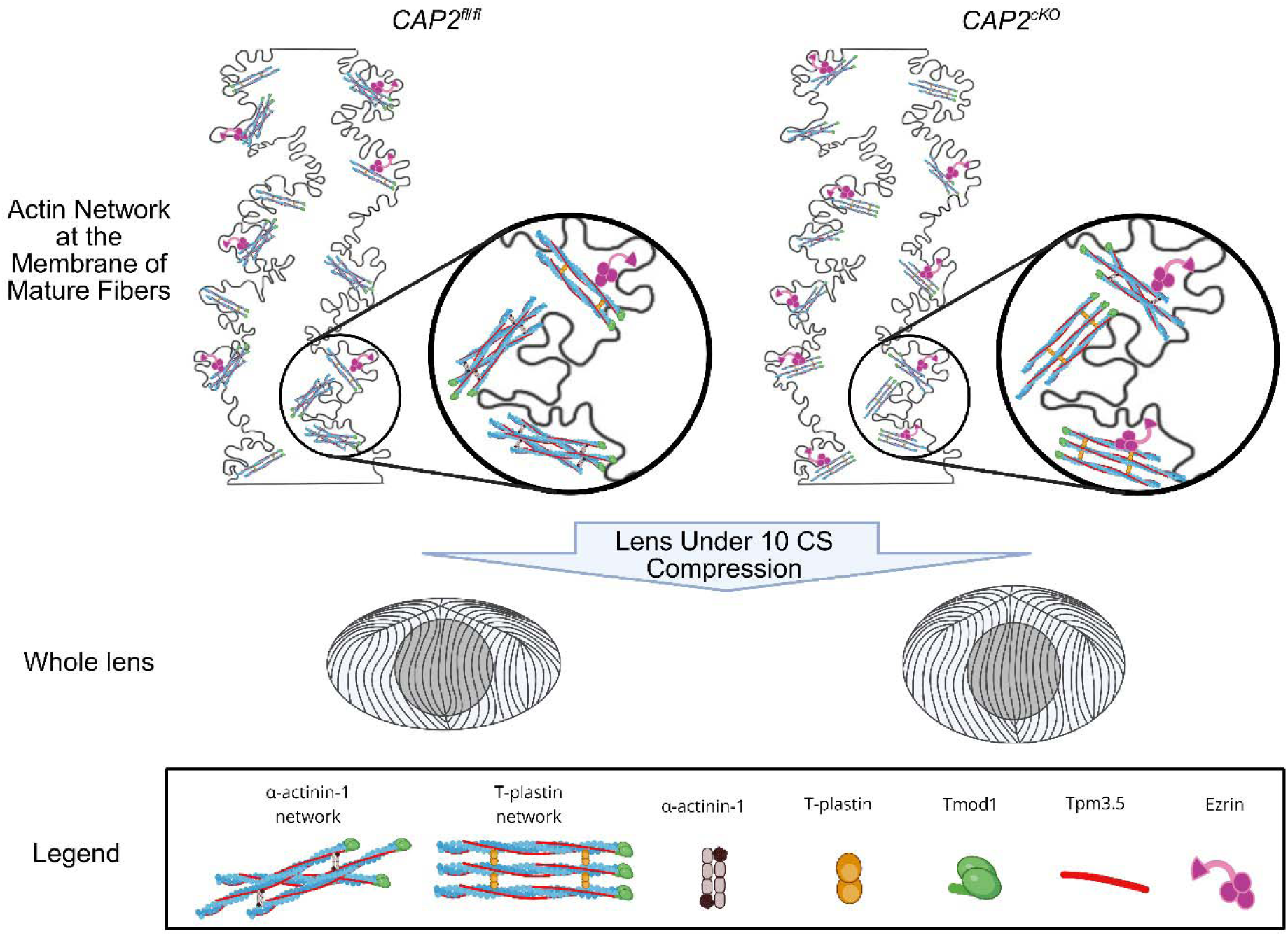
Summary diagram illustrating the effects of *CAP2* deletion on lens biomechanics and membrane-associated F-actin networks. Top: Zoomed-in diagrams of individual mature fiber cells depicting the organization of actin networks at the membrane. Black circles: Close-up views of actin-binding protein interactions at the membrane. Loss of CAP2 leads to enhanced Tmod1 capping and increased association of actin filaments with Tpm3.5 and Ezrin. F-actin filaments are increasingly bundled with T-plastin, and there is a reduction in α-actinin-1–cross-linked filaments in *CAP2^cKO^* lens fibers. The arrow points to lenses from *CAP2^fl/fl^* (left) and *CAP2^cKO^* (right) mice under 10 coverslip (CS) compression, showing decreased axial strain in the *CAP2^cKO^* lens. Bottom black box: Legend of proteins depicted in the diagram. Created in BioRender. Cheheltani, S. (2025) https://BioRender.com/cilm0ic

Using a lens-specific conditional knockout model, we found that CAP2 does not directly affect lens transparency and gross morphology but plays a role in modulating lens stiffness. In contrast, a previous study reported microphthalmia in the global CAP2 knockout mice (Field et al., 2015), likely due to early developmental defects arising from the loss of CAP2 in all ocular tissues, including the retina and optic cup (Harding et al., 2021; Verma and FitzPatrick, 2007). In our model, where CAP2 deletion is restricted to lens epithelial and fiber cells, the lenses displayed a normal shape and did not exhibit any signs of opacities. Yet, they were significantly stiffer under compressive load, suggesting that local cytoskeletal remodeling, rather than large-scale structural disruption, underlies the mechanical phenotype. CAP2’s localization with membrane-associated F-actin in fiber cells points to a role in regulating F-actin turnover near the membrane, where actin network remodeling is likely essential for maintaining biomechanical plasticity.

### CAP2 Regulates Mouse Lens Stiffness Through Local Remodeling of Actin Networks

In comparison to previous studies, which showed that deletion or reduction of actin-stabilizing proteins such as Tmod1 or Tpm3.5 resulted in decreased lens stiffness in the mouse (Cheng et al., 2018; Cheng et al., 2016c; Gokhin et al., 2012; Parreno et al., 2020), the deletion of CAP2 resulted in increased lens stiffness, supporting the idea that CAP2 acts antagonistically to F-actin-stabilizing factors. In cardiomyocytes and other cells, CAP2 enhances the turnover of ADF/cofilin-bound F-actin by targeting filament pointed ends for disassembly (Colpan et al., 2021; Kumar et al., 2016; Makkonen et al., 2013; Quintero-Monzon et al., 2009). In contrast, Tmod and Tpms cap and stabilize actin filament pointed ends, respectively (Cheng et al., 2016c; Kostyukova, 2008; Weber et al., 1994), a relationship shown to oppose CAP-mediated turnover (Colpan et al., 2021; Iwanski et al., 2021). In *CAP2^cKO^*lenses, we observed an increase in both Tmod1 and Tpm3.5 localization within the F-actin-rich membrane protrusions of mature fibers, suggesting that in the absence of CAP2, filaments are stabilized by increased pointed end capping by Tmod1. Additionally, increased Tpm3.5 at the F-actin-rich protrusions of *CAP2^cKO^* lenses supports the view that filament turnover is reduced, likely leading to increased rigidity of the actin networks at the membrane (Fig 8).

Notably, the actin filament stabilization is accompanied by selective remodeling of cross-linked networks, which can further contribute to lens stiffness in the absence of CAP2. α-actinin-1-associated puncta are reduced in cKO lenses, while T-plastin-associated puncta are increased. α-actinin-1 crosslinks actin into loose, contractile arrays of anti-parallel filaments, whereas T-plastin bundles actin into tightly packed, parallel filaments, which form stiffer networks (Bretscher, 1981; Matsudaira, 1994; Winkelman et al., 2016). α-actinin-1 and T-plastin both function as actin-bundling proteins but organize filament networks with distinct geometries and are often mutually exclusive (Cheng et al., 2018; Yamashiro-Matsumura and Matsumura, 1986). This suggests that the shift from α-actinin-1 to T-plastin-associated actin networks could contribute to the increased stiffness by decreasing cytoskeletal flexibility and promoting structural compaction (Yamashiro-Matsumura and Matsumura, 1986). Interestingly, another membrane-associated cytoskeletal component, β2-spectrin, remained unchanged, suggesting that CAP2 loss selectively alters some actin networks without disrupting the underlying membrane-associated scaffold (Cheng et al., 2017; Leterrier and Pullarkat, 2022). This suggests a model in which CAP2 and Tmod1 exist in a local regulatory balance where CAP2 facilitates pointed-end turnover, while Tmod1 caps and stabilizes the ends, as proposed for thin filaments in striated muscle and during cardiac myofibrillogenesis (Iwanski et al., 2021). When CAP2 is removed, this balance shifts toward the stabilization of filaments and the rearrangement of filament networks, ultimately leading to altered fiber cell and lens mechanics, as depicted in Figure 8.

The abundant actin cytoskeleton within fiber cell membrane interdigitations is supported by membrane-cytoskeletal linkers, including components of the ezrin, periplakin, periaxin, and desmoyokin (EPPD) cell junctional complex, which also includes spectrin (Audette et al., 2017; Blankenship et al., 2007; Cheng et al., 2017; Gonen et al., 2004; Lindsey Rose et al., 2006; Rao and Maddala, 2006; Straub et al., 2003). Ezrin, a member of the ERM (ezrin-radixin-moesin) family, links F-actin to the plasma membrane and regulates cell shape and cortical stiffness (Bagchi et al., 2004; Ivetic and Ridley, 2004; Louvet-Vallée, 2000; Rao et al., 2008). In its active form, ezrin undergoes phosphorylation-driven conformational changes that enable its interaction with both F-actin and membrane proteins, facilitating the formation of membrane protrusions, supporting cortical tension, and increasing cytoskeletal stiffness (Viswanatha et al., 2013; Zhang et al., 2020). In *CAP2^cKO^*lenses, we observed increased total ezrin protein levels and enhanced ezrin localization at the membrane of mature fiber cells, particularly at F-actin-rich protrusions. These findings suggest that, in addition to filament stabilization and altered crosslinking, upregulation of ezrin-mediated membrane-actin tethering and its association with increased cortical stiffness may further contribute to the stiffer biomechanical phenotype of CAP2-knockout lenses. Although we detected an increase in total ezrin, further work is needed to determine whether the active form is enriched in the knockout lenses. If so, activated ezrin could further reinforce membrane-cytoskeletal interactions, contributing to increased resistance to deformation.

### Implications for Cap2 functions in cell and tissue mechanics and motility

Beyond the lens, CAP2-mediated actin turnover is broadly required for the formation of dynamic protrusions and efficient migration (Rust et al., 2020). Motility defects are observed in fibroblasts of CAP2-kncokout mice, where the loss of CAP2 results in increased focal adhesions, extended protrusions, and higher F-actin content consistent with stabilization of peripheral actin networks, without changes in total actin. (Kosmas et al., 2015). Similarly, primary keratinocytes from CAP2-knockout mice migrate more slowly in scratch-wound assays, showing a delayed closure of the scratch compared to wild-type cells (Kosmas et al., 2015). In carcinoma models, CAP2 co-localizes with F-actin at the leading edge of lamellipodia, and its depletion impairs lamellipodial extension and motility (Effendi et al., 2013). Conversely, ER-stress-driven upregulation of CAP2 enhances migration and invasion (Yoon et al., 2021). Although these studies did not directly measure other actin-binding proteins, they showed increased F-actin and altered bundling at the cell periphery without changes in total actin (Kosmas et al., 2015), mirroring the redistribution of F-actin stabilizers without overall F-actin changes we observed in the CAP2-knockout lens.

Lens fiber cell paddles and protrusions in the cell-cell interlocking interdigitations are enriched with F-actin that is stabilized by Tpm/Tmod, cross-linked by T-plastin, and anchored to the cell membrane by ERM proteins (Cheng et al., 2018; Gokhin et al., 2012; Straub et al., 2003). They also maintain the fiber-to-fiber cell interactions by cell adhesion molecules, such as cadherins and AQP0 (Bassnett et al., 1999; Wang and Schey, 2011). In lens fibers, CAP2 deletion coincides with increased Tmod1, Tpm3.5, T-plastin, and ezrin, suggesting that CAP2 normally limits actin stabilization at fiber-fiber cell interfaces. In its absence, the actin membrane network becomes more stabilized and rigidly bundled, likely stiffening the paddles and interdigitations and, consequently, the entire lens. To our knowledge, this is the first demonstration that CAP2 regulates cell and tissue biomechanics in a non-muscle system by modulating F-actin-associated proteins, linking molecular control of actin turnover to tissue-level mechanical behavior in an intact vertebrate organ.

## Methods and Materials

### Generation of lens-specific CAP2^cKO^ mice

All animal procedures were conducted in adherence to the ARVO Statement for the Use of Animals in Ophthalmic and Vision Research and performed in accordance with approved animal protocols from the Institutional Animal Care and Use Committee guidelines at the University of Delaware. *CAP2^LoxP^* mouse was obtained from Dr. Jeffrey Field at the University of Pennsylvania (Field et al., 2015). The targeted mouse was created by the European Conditional Mouse Consortium (EUCOMM), which provided the targeting construct and ES clones (*CAP2^tm1a(EUCOMM^*^)^), which Dr. Field crossed with the actin-FLP mouse to excise the FRT sites, *LacZ,* and *neo* cassettes. This process generated the *tm1c* allele, restoring CAP2 expression (Kanuri et al., 2021; Skarnes et al., 2011). This transgenic mouse harbors the CAP2 gene with two LoxP (locus of X-over P1) sites flanking exon 3 (*CAP2^LoxP^* or *CAP2^fl/fl^*) (Field et al., 2015). Exon 3 is crucial for CAP2 function as it encodes the HFD domain responsible for F-actin severing activity. Subsequently, we bred the *CAP2^fl/fl^* mice with *MLR10-Cre* mice (STOCK Tg (Cryaa-Cre)10Mlr/J, Jackson Labs) obtained from Dr. Melinda Duncan at the University of Delaware, resulting in the generation of *CAP2^fl/fl^ ^Tg^ ^(Cryaa-Cre)10Mlr^* (Referred to as *CAP2^cKO^*) transgenic mice with lens-specific CAP2 deletion. The *MLR10-Cre* mouse features a deliberate insertion of a 20-bp Pax6 consensus-binding site within the transgenic αA-crystallin promoter, enabling CRE transgene expression in both lens fiber cells and epithelium (Zhao et al., 2004). CRE expression from these promoters initiates in the lens vesicle at E10.5 in *MLR10-Cre*, leading to the deletion of floxed exon 3 in the lens, thereby knocking out CAP2 in both lens epithelial and fiber cells (Field et al., 2015; Lam et al., 2019). Throughout this study, *CAP2^fl/fl^*mice were used as littermate controls. All mice were genotyped through TransnetYX (Cordova, TN) by semi-quantitative real-time PCR using primer pairs targeting the *CAP2^fl/fl^*, *CAP2^flfl; Tg (Cryaa-Cre)10Mlr^*, and CRE alleles.

### Lens biomechanical testing and morphometrics

Lenses from 8-week-old mice were used for compression assay and morphometric measurements. Freshly dissected lenses were transferred to a custom-made chamber filled with 1X phosphate-buffered saline (PBS) (diluted 1:10 from Gibco PBS (10X), pH 7.4, 70011044, Thermo Fisher Scientific), and top-view images were acquired using an Olympus SZ11 dissecting microscope attached to a digital camera. To test the biomechanical properties of the lens, we conducted a coverslip compression assay as previously described (Cheng et al., 2016b). A series of ten 18×18 glass coverslips (∼114.8 mg each) (12542A, Fisherbrand, Pittsburgh, PA, USA) was sequentially placed on top of freshly dissected lenses, positioned in a divot inside the chamber. Side-view images of the lenses were captured using a 45-degree angled mirror. The equatorial and axial diameters of the lenses before and after each loading step were measured using FIJI software. Axial and equatorial strains were calculated from: ε = (d - d0) / d0, where ε is strain, d denotes the axial or equatorial diameter at a given load, and d0 represents the corresponding axial or equatorial diameter at zero load (Cheng et al., 2016a; Fudge et al., 2011; Gokhin et al., 2012). Lens volume was calculated from: volume = 4/3 × π × r_E_² × r_A_, where r_E_ is the equatorial radius and r_A_ is the axial radius (Sindhu Kumari et al., 2015). The lens aspect ratio was determined by dividing the equatorial diameter by the axial diameter. To evaluate the morphology of the lens nucleus, we meticulously removed peripheral fiber cells by gently rolling the lens between gloved fingertips, revealing a densely compacted central region known as the lens nucleus. Nuclear volume was calculated from volume = 4/3 × π × r_N_³, where r_N_ is the radius of the lens nucleus. The nuclear fraction is the ratio of the nuclear volume to the total lens volume (Gokhin et al., 2012).

### RNA isolation and RT-qPCR

Two lenses of each mouse were pooled and homogenized in 100 μL of TRIzol reagent (15596026, Thermo Fisher Scientific) for RNA isolation. The amount of RNA was measured by A_260_, and an equal amount of RNA from each lens sample was used for cDNA synthesis. Reverse transcription and cDNA synthesis were performed using the Superscript^TM^ III First-Strand Synthesis System kit (18080051, Thermo Fisher Scientific). RT-qPCR was performed on an equal amount of cDNA from each sample in a 20 µl volume containing TaqMan™ Fast Advanced Master Mix for qPCR (4444557, Thermo Fisher Scientific), TaqMan probes for Cap1 (Mm00482950_m1, Thermo Fisher Scientific), Cap2 spanning exons 2 and 3 (Mm00482639_m1, Thermo Fisher Scientific), and GAPDH (Mm99999915_g1, Thermo Fisher Scientific) as the housekeeping gene for normalization. Three lens pairs from each genotype were used for each experiment.

### Western blotting

Western blots were performed on lenses isolated from 8-week-old mice, as previously described (Cheng et al., 2018). Freshly dissected lenses were stored at -80°C until homogenization. Two lenses of each mouse were pooled into a single protein sample. Lens pairs were homogenized on ice in 250 µL of lens homogenization buffer (20 mM Tris-HCl pH 7.4 at 4°C, 100 mM NaCl, 1 mM MgCl_2_, 2 mM EGTA and 10 mM NaF with 1 mM DTT), 1:100 Protease Inhibitor Cocktail (P8430, Sigma-Aldrich) and 1:1000 Phosphatase Inhibitor (78420, Thermo Fisher Scientific) per 10 mg of lens wet weight. The lysates were then diluted 1:1 with 2× Laemmli sample buffer (1610737, Bio-Rad Laboratories, Hercules, CA, USA). Samples were briefly sonicated with a Q55 Sonicator (Qsonica, Newtown, CT, USA) and boiled for 5 minutes. Proteins were separated by electrophoresis on 4-20% linear gradient SDS-PAGE mini-gels (XP04205BOX, Thermo Fisher Scientific) and transferred to nitrocellulose membranes (10600011, Amersham Protran, Slough, UK) at 150V in 1× transfer buffer (25 mM Tris, 192 mM glycine in ddH_2_O) with 20% methanol in a trans-blot tank (Bio-Rad) at 4°C for 1 hour. Membranes were stained with Ponceau S (09189, Fluka BioChemica, Mexico City, Mexico), and gently washed with ddH_2_O. The blots were scanned with a Bio-Rad ChemiDoc MP to reveal total protein levels in each lane and blocked with 5% BSA in 1× PBS for 1 hour at room temperature. The blots were then incubated with primary antibodies diluted in 5% BSA + 0.1% Triton X-100 in PBS overnight at 4°C with gentle rocking.

The primary antibodies used for western blotting were anti-actin (C4, 1:20,000, Millipore, Burlington, MA), anti-CAP2 (15865-1-AP, 1:1000 Proteintech), anti-α-actinin-1 (non-sarcomeric, Actn1, A5044, 1:1000, Sigma-Aldrich), anti-T-plastin (ab137585, 1:1000, Abcam), anti-Tmod1 (NBP2-00955, 1:1000, Novus Biologicals), anti-Tpm3.5 (CH1, 0.5 ug/ml, Developmental Studies Hybridoma Bank, Iowa City, IA), anti-ezrin (E8897, 1:1000, Sigma-Aldrich), and anti-GAPDH (NB300-221, 1:1000, Novus Biologicals). The secondary antibodies (1:20,000) were IRDye-800CW-conjugated goat anti-rabbit-IgG (926-32211, LI-COR, Lincoln, NE), IRDye-680LT-conjugated goat anti-mouse-IgG (926-68020, LI-COR, Lincoln, NE), or IRDye-680RD-conjugated goat anti-mouse IgM (µ chain specific) (925-68180, LI-COR, Lincoln, NE). After incubation with secondary antibodies, the blots were washed with 1X Tris-buffered saline with 0.1% Tween® 20 detergent (TBST) (4 × 5 minutes per wash). Antibodies and other reagents are also listed in Table S1. The band intensities of the blot were quantified using FIJI with background subtraction and normalized to total protein (Ponceau S staining) or GAPDH. Three lens pairs from each genotype were used for each experiment.

### Immunostaining of frozen sections

To prepare frozen sections of the eye, a small opening was made at the corneal-scleral junction of freshly enucleated eyeballs to allow penetration of fixative (Cheng et al., 2016c). Eyeballs were fixed for 4 hours at 4°C in 1% paraformaldehyde (PFA) in PBS, prepared fresh from a 16% stock (15710, Electron Microscopy Sciences, Hatfield, Pennsylvania). Samples were subsequently washed in cold PBS 3 times and cryoprotected in 30% sucrose for ∼2 hours, until sinking, and then embedded in OCT medium (Sakura Finetek, Torrance, CA) in cross or sagittal-section orientation in blocks. Frozen sections (12 μm thick) were collected with a Leica CM3050 cryostat and stored at -20°C. Sections were briefly rehydrated with PBST (1× PBS, 0.01% Triton X-100) and permeabilized and blocked in blocking buffer (3% BSA, 3% goat serum, and 0.3% Triton X-100 in 1× PBS) for 1 hour before incubation with the primary antibody. Lens sections were labeled with primary antibody 1:100 in blocking buffer overnight at 4°C, washed in PBST (3 × 5 minutes) and labeled for 90 minutes with secondary antibody along with rhodamine phalloidin (R415, 220 nM, Thermo Fisher Scientific), WGA (Alexa Fluor™ 488 Conjugate, W11261, 1:500, Life Technologies) and Hoechst 33342 (62249, 1:1000 dilution, Thermo Fisher Scientific). Secondary antibodies IgG (1:200 dilution) were Alexa-Fluor-647-conjugated goat anti-mouse IgG (A21236, Thermo Fisher Scientific) or Alexa-Fluor-647-conjugated goat anti-rabbit. Sections were washed in PBST (4 × 5 minutes) and mounted with ProLong® Gold antifade reagent (P36934, Thermo Fisher Scientific), and sealed with clear nail polish. Imaging was performed using a Zeiss LSM880 confocal microscope (20× objective, NA 0.8 or 63× oil immersion objective, NA 1.4).

### Immunostaining of single fiber cells

Single fiber cells were prepared as previously described (Vu and Cheng, 2023). Briefly, lenses were dissected from freshly enucleated eyeballs, and the lens capsule was removed using sharp tweezers. Decapsulated lenses were fixed in 1% PFA overnight at 4°C with gentle rocking. Fiber masses were then cut into quarters in sagittal orientation using a scalpel and post-fixed in 1% PFA for 15 minutes with gentle rocking at room temperature. Lens quarters were then washed in 1×PBS (2 × 5 minutes each) and blocked for 1 hour in PBS containing 3% BSA, 3% goat serum, and 0.3% Triton X-100. Quarters were then incubated overnight at 4°C with gentle rocking in a 1:50 dilution of the primary antibody. After washing the lens quarters with PBST (3 × 10 minutes), they were labeled with secondary antibodies (1:100), rhodamine phalloidin (1:100), and WGA (1:100) for 3 hours at room temperature. After incubation with secondary antibodies, the lens quarters were washed again (3 × 10 minutes). Then, fiber cells were separated from each other using two tweezers on a drop of ProLong Gold antifade reagent. A coverslip was then sealed onto each slide. Z-stacks of single fiber cells were acquired at a digital zoom of 3.0 and a z-step of 0.17µm using a Zeiss LSM880 microscope (63× oil immersion objective, NA 1.4, AiryScan Super-resolution).

### Whole mount staining and capsule thickness measurement

In this procedure, whole fixed lenses were fluorescently labeled with rhodamine-phalloidin (1:300), WGA (1:250), and Hoechst 33342 (1:500), as previously described (Emin et al., 2024). Each lens was positioned on a glass-bottom dish with its anterior surface facing the inverted microscope objective to visualize the anterior capsule. Z-stacks of the capsule thickness were acquired with digital zoom of 1.0 with z-step size of 0.3µm using a Zeiss LSM880 confocal microscope with a 40× oil immersion objective, NA 1.3. Images were processed in Zeiss Zen Software (Blue 3.7) at the X-Z plane, and capsule thickness was measured as described previously(Cheheltani et al., 2025a; Parreno et al., 2018). Briefly, the distance between the top surface of the capsule stained by WGA and the membrane of the underlying actin cytoskeleton stained by rhodamine phalloidin was measured using line scan analysis in FIJI (Fig. S4A-C).

### Image analysis

To assess fiber cell organization in *CAP2^cKO^* and *CAP2^fl/fl^* control lenses, we quantified regions of disordered fiber cell packing in equatorial cryosections labeled with F-actin, as previously described (Islam et al., 2024; Islam et al., 2023). Briefly, single optical sections captured at 20× magnification were analyzed in FIJI by manually outlining areas of disrupted fiber organization. The area of each disordered region was measured, and the percent disorder was calculated by dividing the total disordered area by the area of the region of interest. Three sections from at least three different mice per genotype were analyzed for this quantification

To determine the fluorescent intensity in lens cryosections, the split channel function in the FIJI software was used to separate the red and green channels. The same area of the fiber cell was then outlined, and the mean fluorescent intensity was measured. The mean fluorescent intensity of rhodamine phalloidin was normalized to WGA fluorescent intensity. To quantify the fluorescent intensity of ABPs in lens cryosections, the signal intensity of the protein of interest was normalized to rhodamine phalloidin. Six to seven different sections from three mice per genotype were used.

To determine the fluorescent intensity of individual ABPs in single fiber cells, Z-stack confocal images were further analyzed in Zen 3.7 microscopy software (Zeiss). Maximum intensity projections from Z-stacks were exported into FIJI, and the mean fluorescent intensity in the membrane protrusions of mature fiber cells was measured by outlining the bright rhodamine-phalloidin-stained protrusion area of each single fiber cell with the FIJI freehand tool (Fig. S7). For each cell, the mean fluorescence intensity of each protein in membrane protrusions was divided by the mean rhodamine–phalloidin intensity measured for that same cell, yielding a normalized intensity. Data represent means determined for 20-30 single fiber cells from 3 different mice per genotype.

### Statistical Analysis

Statistical significance was determined using the two-tailed Student *t*-tests or One-way ANOVA, and mean and s.d. were calculated and plotted using GraphPad Prism 9. A p-value less than or equal to 0.05 was considered statistically significant.

## Acknowledgements

The authors thank the University of Delaware animal facility and Dr. Melinda Duncan for providing us with a breeding pair of the MLR10-Cre mice, as well as Dr. Salil Lachke for sharing unpublished data on the phenotype of the CAP2 global KO mouse lens, leading us to study the lens-specific CAP2-knockout mouse lens.

## Competing interests

The authors declare no competing or financial interests.

## Funding

This work was supported by a grant from the National Eye Institute at the National Institutes of Health [R01 EY017724 to V.M.F]. This research also benefited from the BioStore data management services of the Delaware Biotechnology Institute and Center for Bioinformatics and Computational Biology at the University of Delaware (RRID: SCR_017696), supported by NIH (NIGMS S10OD028725) and DE-INBRE (NIH/NIGMS P20GM103446).

## Notes

### Competing Interest Statement

The authors have declared no competing interest.

