## Supplementary material for "Cyclase associated actin cytoskeleton regulatory protein 2 (CAP2) organizes the actin cytoskeleton to influence the biomechanical properties of the ocular lens"

Table of Materials

| **Material** | **Manufacturer** | **Catalog #** |
|---|---|---|
| TRIzol™ Reagent | Thermo Fisher Scientific | 15596026 |
| Tris base ((hydroxymethyl)aminomethane) | Thermo Fisher Scientific | BP152-500 |
| HCl (hydrochloric acid) | Sigma-Aldrich | 320331 |
| NaCl (sodium chloride) | Thermo Fisher Scientific | S271-500 |
| MgCl2 (magnesium chloride) | Thermo Fisher Scientific | BP214-500 |
| EGTA (ethylene glycol-bis (β-aminoethyl ether) | bioWORLD | 40520008-1 |
| NaF (sodium fluoride) | Spectrum | S0167 |
| DTT (Dithiothreitol) | Thermo Fisher Scientific | R0861 |
| Glycine | MP Biomedicals | 808822 |
| SDS (sodium dodecyl sulfate) | Sigma-Aldrich | L3771 |
| Methanol | Fisher Scientific | A454-4 |
| Protease Inhibitor Cocktail | Sigma-Aldrich | P8430 |
| Phosphatase Inhibitor | Thermo Fisher Scientific | 78420 |
| 2X Laemmli sample buffer | BIO-RAD | 1610737 |
| Ponceau S solution 0.1% (w/v) in 5% acetic acid | Fluka BioChemica | 9189 |
| 4–20% linear gradient SDS-PAGE mini-gels | Thermo Fisher Scientific | XP04205BOX |
| Nitrocellulose membrane | Amersham Protran | 10600011 |
| Transfer blot tank | BIO-RAD | 1703930 |
| Bovine Serum Albumin (BSA) Fraction V ≥98% Purity | Genesee Scientific | 25-529 |
| 10X phosphate-buffered saline (PBS) | Thermo Fisher Scientific | 70011044 |
| Triton™ X-100 | Thermo Fisher Scientific | 28314 |
| IRDye-680LT-conjugated goat anti-mouse-IgG | LI-COR | 926-68020 |
| IRDye-800CW-conjugated goat anti-rabbit-IgG | LI-COR | 926-32211 |
| IRDye 680RD Goat anti-Mouse IgM (µ chain specific) | LI-COR | 925-68180 |
| Paraformaldehyde (16%) | Electron Microscopy Sciences | 15710 |
| Goat Serum | Invitrogen | A11008 |
| Q55 Sonicator Ultrasonic Processor | Qsonica | Mfr # Q55-220 |
| ChemiDoc MP Imaging System | BIO-RAD | 12003154 |
| Olympus SZ11 dissecting microscope | Olympus | 8F20022 |

| | | |
|---|---|---|
| Swiftcam 20 Megapixel Camera | Amazon | SC2003 |
| FluoroDish cell culture dishes | WPI | FD35-100 |
| Sucrose | Sigma-Aldrich | 84097-1KG |
| Tissue-Tek OCT Compound | Sakura Finetek | 4583 |
| Leica CM3050 cryostat | Leica Biosystems | 14903050S03 |
| CrystalCruz® Adhesive Micro Slides | CrystalCruz | sc-363560 |
| Coverslip #1.5 18 x 18 mm (Lens compression assay) | Thermo Fisher Scientific | 12-542-AP |
| Coverslip #11/2 size 22x 40mm | Electron Microscopy Sciences | 72204-03 |
| ProLong® Gold antifade reagent | Thermo Fisher Scientific | P36934 |
| Rhodamine phalloidin | Thermo Fisher Scientific | R415 |
| Wheat Germ Agglutinin (WGA) conjugated to Alexa Fluor-488 | Thermo Fisher Scientific | W11262 |
| Hoechst 33342 | Thermo Fisher Scientific | H3570 |
| Alexa-Fluor-647-conjugated goat anti-rabbit-IgG | Thermo Fisher Scientific | A-21245 |
| Alexa-Fluor-488-conjugated goat anti-mouse-IgG | Thermo Fisher Scientific | A11001 |
| Alexa-Fluor-488-conjugated goat anti-rabbit-IgG | Thermo Fisher Scientific | A11008 |
| Alexa-Fluor-647-conjugated goat anti-mouse-IgG | Thermo Fisher Scientific | A21236 |
| Actin (C4) primary antibody | Millipore | MAB1501 |
| Rabbit Polyclonal Anti-CAP2 antibody | Proteintech | 15865-1-AP |
| Mouse Monoclonal Anti-Tmod1 antibody | Novus Biologicals | NBP2-00955 |
| Mouse monoclonal Anti-Tpm3.5 antibody | DSHB | CH1 |
| Rabbit Polyclonal Anti-T-Plastin antibody | Abcam | ab137585 |
| Mouse monoclonal Anti-α-actinin-1 antibody | Sigma | A5044 |
| Mouse monoclonal Anti-Ezrin antibody | Sigma | E8897 |
| Mouse monoclonal Anti-β2-spectrin antibody | BD Transduction Laboratories | 612563 |
| Mouse monoclonal Anti-GAPDH antibody | Novus Biologicals | NB300-221 |

**Table S1. List of materials, biochemicals, antibodies, and dyes.**

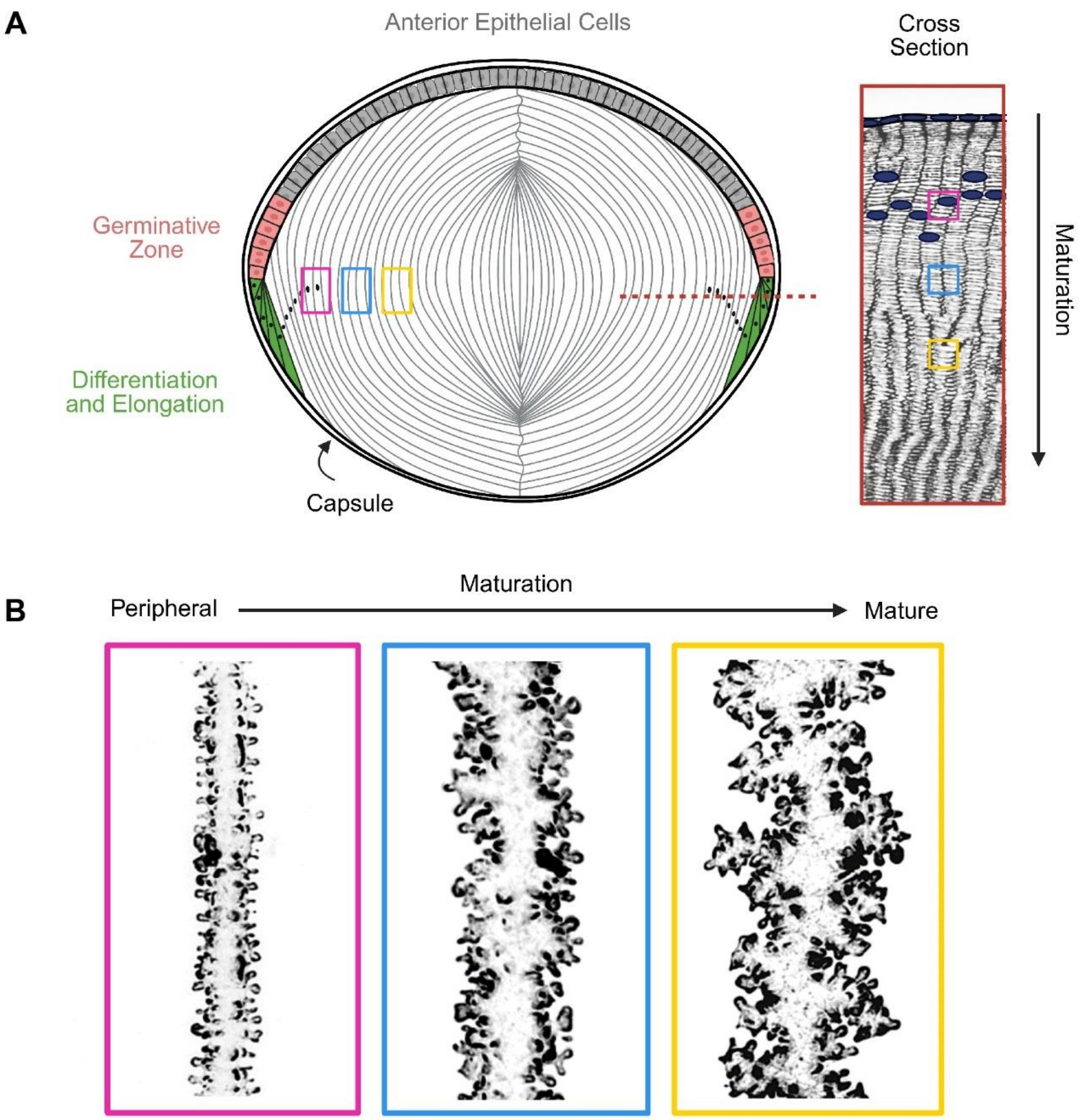


**Figure S1. Lens anatomy diagrams.** (A) A sagittal section of the lens shows an anterior region of epithelial cells (gray) surrounded by a bulk mass of lens fiber cells (white). The whole lens is encapsulated in a basement membrane called the capsule (Created in BioRender. Cheheltani, S. (2025) https://BioRender.com/j3cx4b3). The red dotted line shows a region where the cross-section in the red box is associated with. (B) Representative image of fiber cells at different stages of maturation. Peripheral fiber cells have small protrusions along their membrane (pink box). During maturation, cortical fiber cells develop larger protrusions along the short sides (blue box), and mature fiber cells from the organelle-free zone have large paddle domains decorated by small protrusions (yellow box).

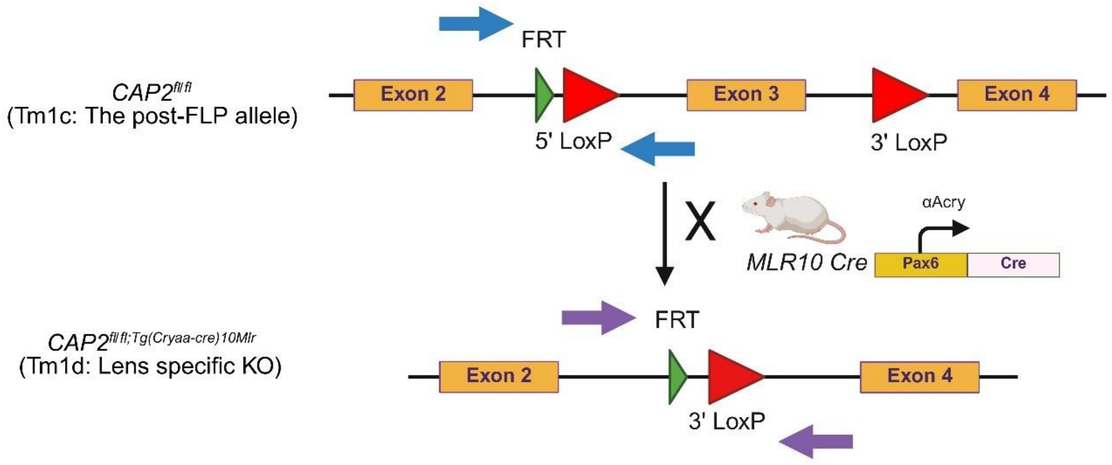


**Figure S2. Diagram of the strategy to create the lens-specific CAP2 KO mouse.** Mice homozygotes for the *CAP2*$^{fl/fl}$ allele were crossed with *MLR10-Cre* transgenic mice to generate lens-specific *CAP2* KO mice. The blue arrows indicate the location of the reverse and forward primers for genotyping the *CAP2*$^{fl/fl}$ allele, which is used as a littermate control, and the purple arrows indicate the location of the primers used to genotype the *CAP2*$^{cKO}$ allele (Created in BioRender. Cheheltani, S. (2025) https://BioRender.com/9kl05s5). The primer design and genotyping were performed by TransnetYX using real-time PCR.

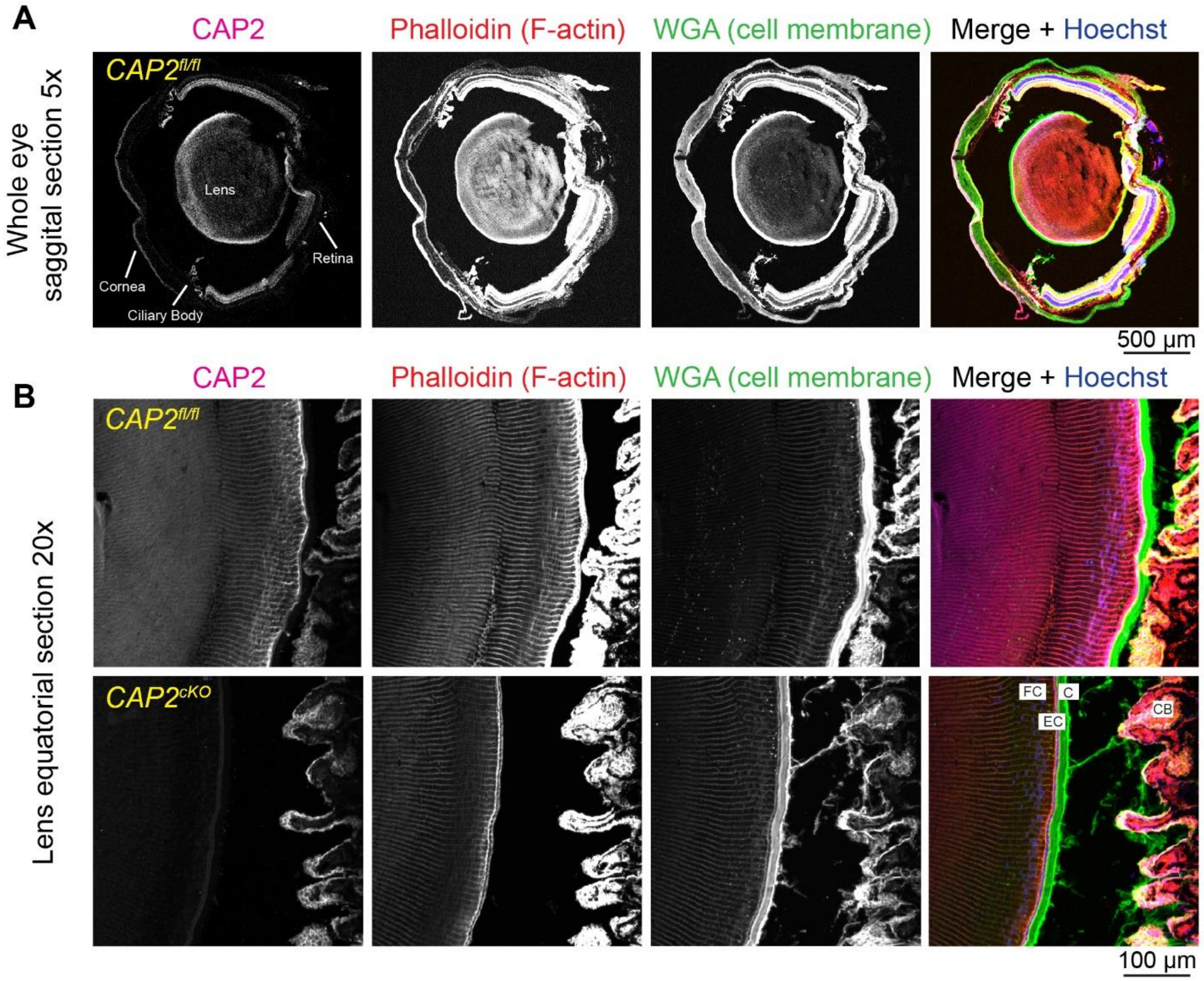


**Figure S3. CAP2 is expressed in multiple ocular tissues but is absent in the lens of *CAP2$^{cKO}$* mice.** (A) Representative 5x confocal images of a sagittal section from an 8-week-old *CAP2$^{fl/fl}$* and *CAP2$^{cKO}$* mouse eye immunostained for CAP2 (magenta), F-actin (red), membrane (green), and nuclei (blue). In control eyes (*CAP2$^{fl/fl}$*), CAP2 is detected in the lens, cornea, retina, and ciliary body. (B) Higher magnification (20×) images of the lens equatorial region show robust CAP2 expression in the lens fiber cells and ciliary body of the control mice, but complete loss of CAP2 signal in the lens of *CAP2$^{cKO}$* mice, confirming tissue-specific deletion. Fiber Cell (FC), Epithelial Cell (EC), Capsule (C), and Ciliary Body (CB). Scale bars, 500 µm (A), 50 µm (B).

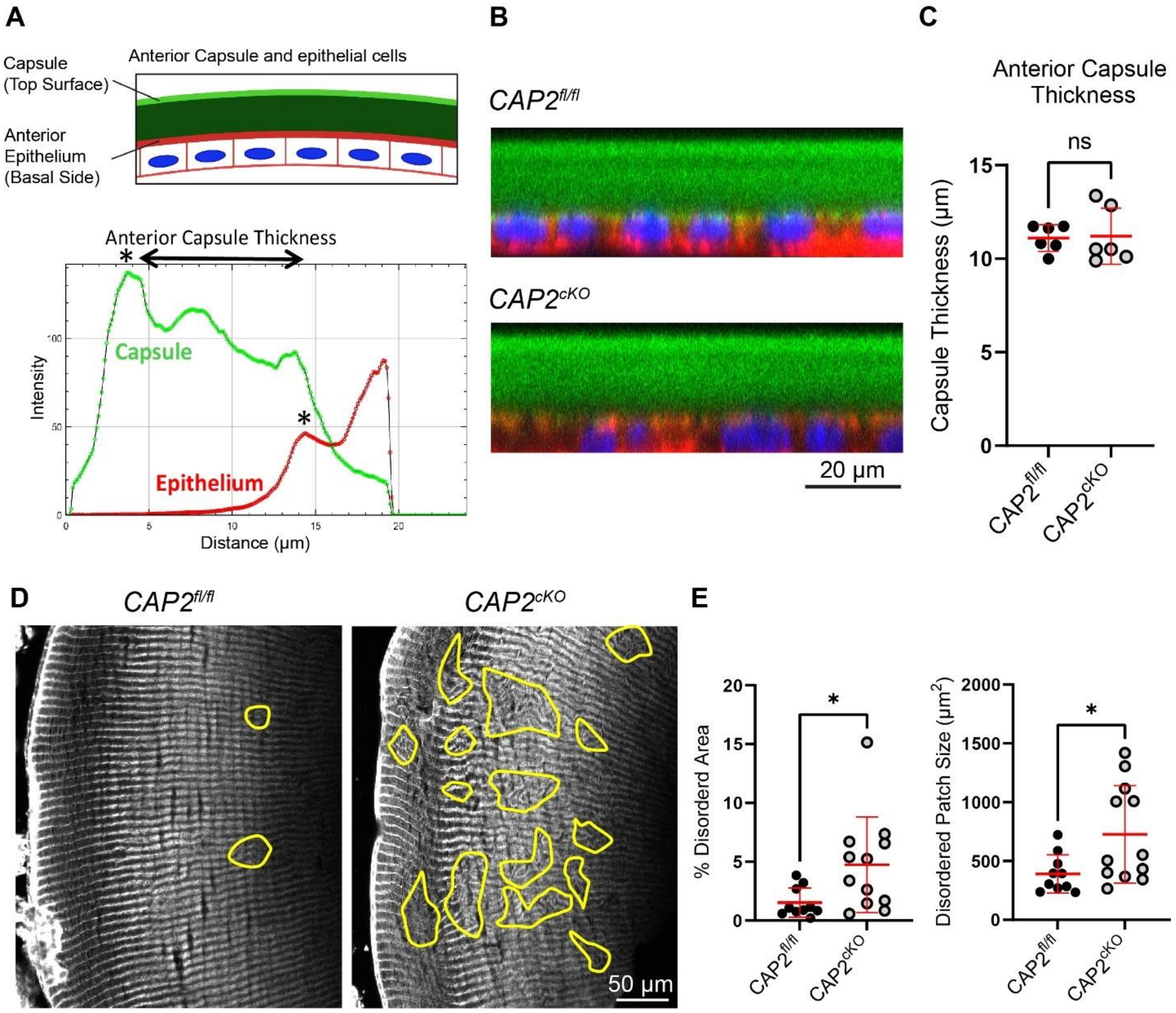


**Figure S4. *$CAP2^{cKO}$* lenses have normal capsule thickness but mild disorganization of fiber cell packing.** (A) Schematic of the anterior lens capsule and epithelium with a representative intensity profile from a line scan across the capsule (green, WGA) and basal F-actin (red, phalloidin) (B) Confocal X-Z reconstruction of lenses from 6-week-old *$CAP2^{fl/fl}$* and *$CAP2^{cKO}$* mice stained with WGA (green), phalloidin(red), and Hoechst (blue) (C) Quantification shows no significant difference in anterior capsule thickness between genotypes (n = 6 lens from 6 mice per genotype). (D) Equatorial sections of lenses stained with rhodamine-phalloidin reveal disorganized regions of fiber cell packing in *$CAP2^{cKO}$* lenses, outlined in yellow. (**E)** Quantification of disordered patch size and percent disordered area indicates slightly higher disorganization in *$CAP2^{cKO}$* lenses compared to controls. Scale bar, 20 µm (B), 50 µm (D). Plots reflect the mean ± s.d. of 2-3 independent immunostained sections from five different mice per genotype. *$P < 0.05$.

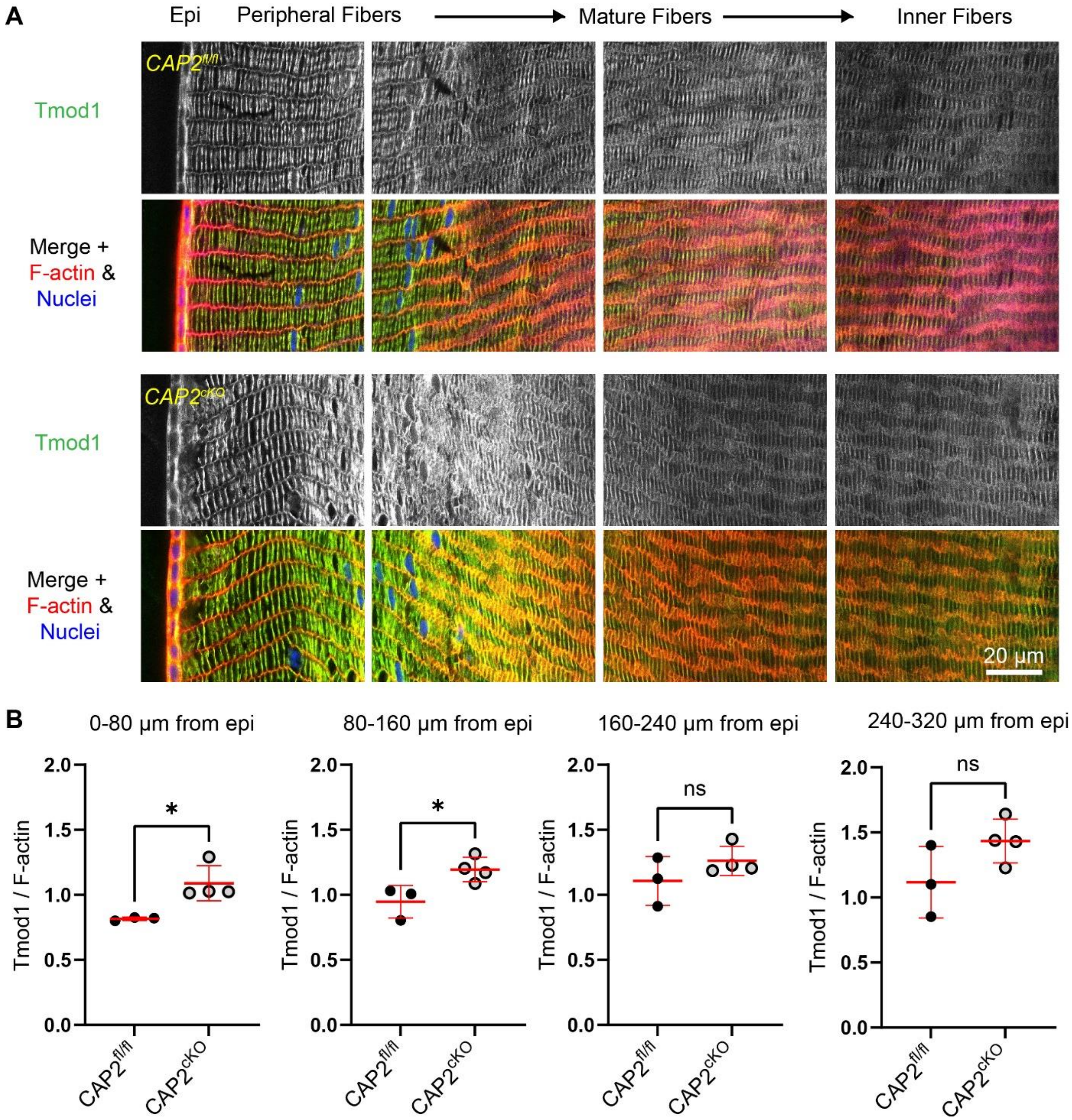


**Figure S5. Increased Tmod1 levels in the outer fiber cell regions of *CAP2*$^{cKO}$ lenses compared to controls.** (A) Representative equatorial cryosections from *CAP2*$^{fl/fl}$ and *CAP2*$^{cKO}$ mouse lenses stained for Tmod1 (green), F-actin (red), and nuclei (blue). Images are shown across four regions spanning from the epithelial layer through peripheral, mature, and inner fiber cells. Tmod1 signal appears more intense in the outer fiber cell regions of *CAP2*$^{cKO}$ lenses. Scale bar, 20 µm. (B) Quantification of Tmod1 fluorescence intensity normalized to F-actin in four successive 80-µm regions extending from the lens epithelium. Tmod1 levels were significantly elevated in the 0-80 µm and 80-160 µm regions of *CAP2*$^{cKO}$ lenses compared to controls, while no significant differences were observed in deeper regions (160-240 µm and 240-320 µm). Data are presented as mean ± s.d. from 3-4 biological replicates per group. *$P$ < 0.05

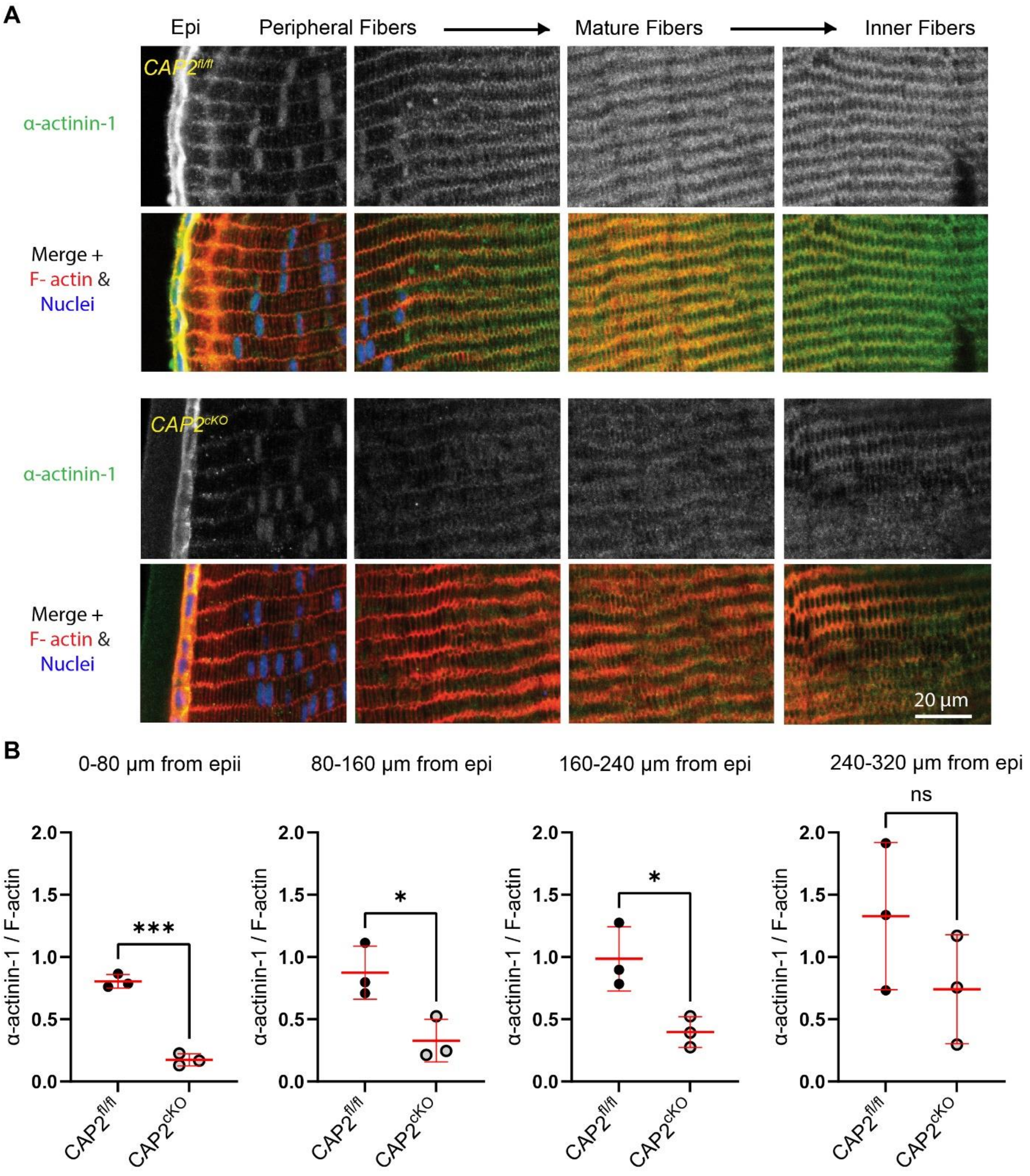


**Figure S6. α-actinin-1 levels are significantly reduced in fiber cells of *CAP2*$^{cKO}$ lenses compared to controls.** (A) Representative equatorial cryosections of *CAP2*$^{fl/fl}$ and *CAP2*$^{cKO}$ lenses stained for α-actinin-1 (green), F-actin (red), and nuclei (blue). Sections span from the epithelium through peripheral, mature, and inner fiber cells. α-actinin-1 signal intensity is visibly decreased in CAP2$^{cKO}$ lenses, particularly in the outer and intermediate fiber regions. Scale bar, 20 µm. (B) Quantification of α-actinin-1 fluorescence intensity normalized to F-actin in four successive 80-µm regions from the epithelium. *CAP2*$^{cKO}$ lenses showed significantly reduced α-actinin-1 levels in the 0-80 µm, 80-160 µm, and 160-240 µm regions compared to controls. No significant difference was observed in the deepest region (240-320 µm). Data represent mean ± s.d. from 3 biological replicates per group. $^*P < 0.05$; $^{***}P < 0.001$.

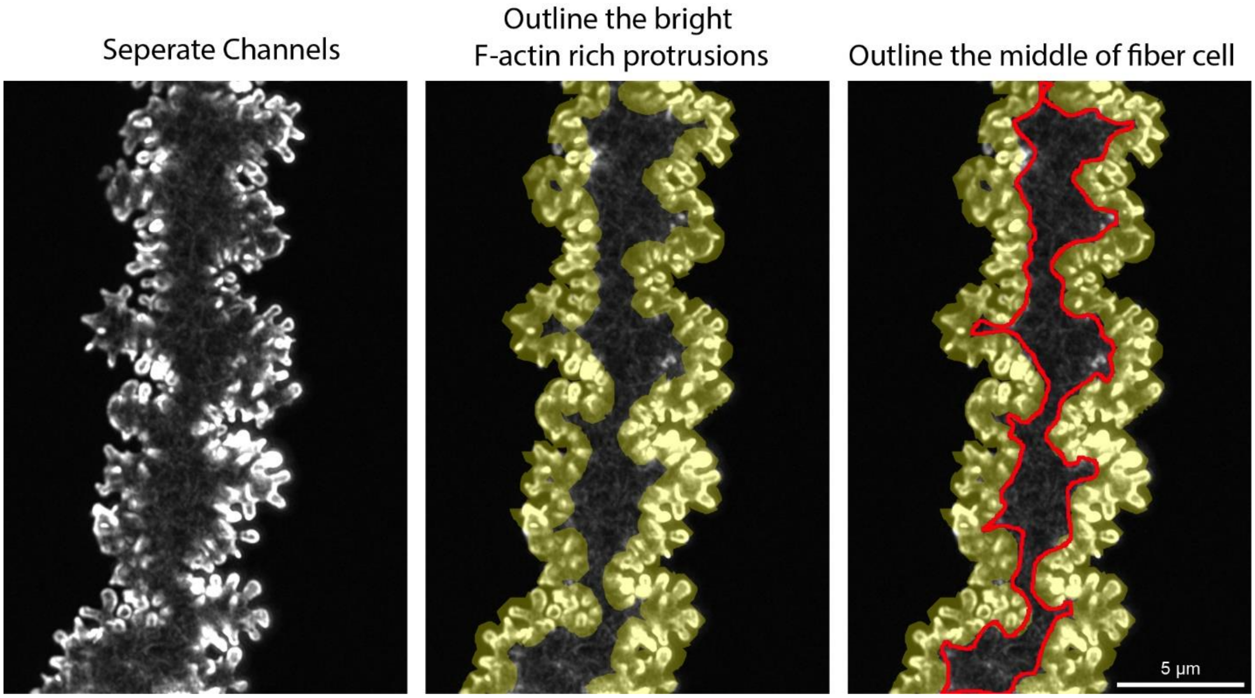


**Figure S7. Method for quantifying fluorescence intensity for the protrusions and cytoplasmic regions of mature lens fiber cells.** Representative image of a mature single fiber cell stained with rhodamine phalloidin for F-actin. Left: F-actin channel shown in grayscale. Middle: The bright F-actin-rich protrusion areas were outlined using the freehand line tool in FIJI. Right: The remaining cytoplasmic area was traced using a freehand selection tool (red outline) to identify the central region of the same fiber. Fluorescence intensity was measured separately for the protrusion and the cell body, and the values were summed for total intensity quantification. Scale bar, 5µm.
